# Expanding the Ligandable Chemical Space of OTUB1 through Discovery of a Four-Membered-Ring Recruiter Chemotype

**DOI:** 10.64898/2026.08.26.747398

**Authors:** Qiong Wu, Xiangyang Song, Li Chen, Hiroyuki Inuzuki, Janell Atkins, Yihang Qi, Yan Xiong, Wenyi Wei, Jian Jin

## Abstract

Deubiquitinase-targeting chimeras (DUBTACs) have emerged as a promising strategy for targeted protein stabilization, but their broader application remains limited by the scarcity of ligandable deubiquitinase recruiters. Here, we report a previously unexplored four-membered-ring OTUB1 recruiter chemotype. Through systematic structure–activity relationship studies, we identified compound **21** (MS2159) as a potent and selective covalent OTUB1 ligand. Biochemical and intact protein mass spectrometric analyses demonstrated that MS2159 selectively engages the non-catalytic C23 residue of OTUB1, shows minimal reactivity toward other tested proteins, and preserves OTUB1 deubiquitinase activity. Conjugation of MS2159 with the CFTR ligand lumacaftor yielded compound **25** (MS2134), which effectively stabilized ΔF508-CFTR. Collectively, these findings establish a new OTUB1 recruiter scaffold, expand the ligandable chemical space of OTUB1, and provide additional opportunities for developing next-generation DUBTACs.

## INTRODUCTION

Targeted protein degradation (TPD) technologies, including proteolysis-targeting chimeras (PROTACs) and molecular glues, have transformed modern drug discovery by enabling selective elimination of disease-associated proteins. By harnessing cellular protein quality-control machinery, these approaches have expanded the druggable proteome beyond the limits of conventional occupancy-driven pharmacology.^1–3^ However, many human diseases arise from insufficient protein abundance, loss-of-function mutations, protein misfolding, or accelerated protein degradation rather than excessive protein activity.^4, 5^ Consequently, restoring protein abundance and function, rather than inhibiting or eliminating the target protein, often represents a more desirable therapeutic strategy in these disease settings. Accordingly, targeted protein stabilization (TPS) has emerged as a complementary induced-proximity modality for restoring protein abundance and function.^6^

Among recently developed TPS technologies, deubiquitinase-targeting chimeras (DUBTACs) represent a particularly attractive induced-proximity strategy for restoring protein abundance. DUBTACs stabilize proteins by recruiting deubiquitinases (DUBs) to proteins of interest, thereby removing degradative ubiquitin chains and rescuing target proteins from proteasomal degradation.^7, 8^ Among the deubiquitinases explored for TPS applications,^9–17^ OTUB1 has emerged as one of the most promising recruiters because it selectively cleaves K48-linked polyubiquitin chains and contains a ligandable non-catalytic cysteine residue (C23) that can be covalently engaged without disrupting enzymatic activity.^18, 19^ These features have established OTUB1 as the most extensively investigated recruiter for DUBTAC development.^9, 10, 12, 15–17^

The successful application of OTUB1-based DUBTACs has stimulated growing interest in the discovery of OTUB1 recruiters. EN523 (**Figure 1**) was the first covalent ligand reported to engage the non-catalytic C23 residue of OTUB1, and subsequent studies identified additional recruiter chemotypes, including MS5105 and our recently reported recruiter MS8572 (**Figure 1**).^9, 12, 20^ Despite these advances, the number of available OTUB1 recruiter scaffolds remains limited. In contrast, the rapid advancement of TPD technologies has been facilitated by the availability of multiple recruiter chemotypes, enabling optimization of target engagement, selectivity, linker architecture, and pharmacological properties.^1, 21^ The emergence of structurally distinct OTUB1 recruiters suggests that productive OTUB1 engagement can be achieved through multiple chemical architectures. Continued expansion of recruiter diversity may therefore provide new opportunities for DUBTAC development and facilitate a deeper understanding of the molecular determinants governing productive OTUB1 recruitment.

**Figure 1.**
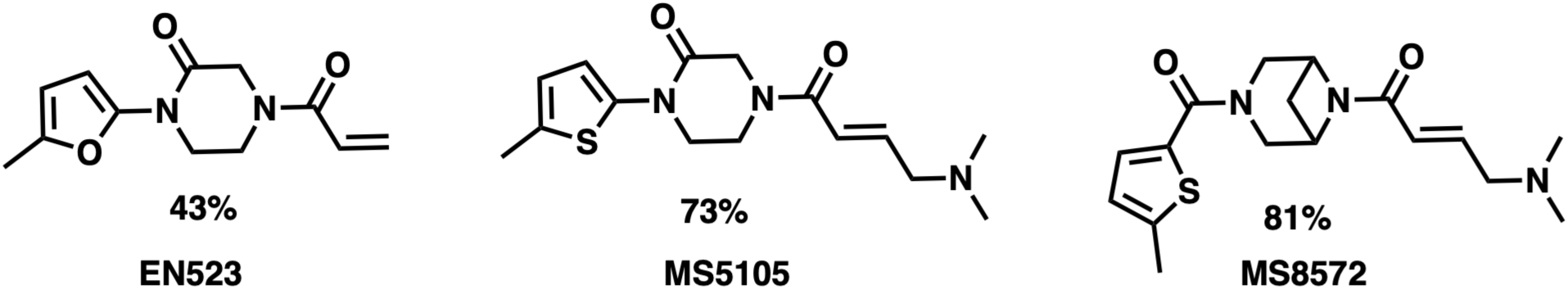
**Reported OTUB1 binders and their rates of covalent OTUB1 modification.**

Here, we sought to expand the repertoire of OTUB1 recruiters through systematic scaffold diversification. Through exploration of heterocyclic scaffold architectures followed by optimization of the covalent warhead and aromatic substituent regions, we identified a previously unrecognized four-membered-ring OTUB1 recruiter chemotype. This effort led to the discovery of compound **21** (MS2159), a potent and selective covalent ligand that efficiently engages the non-catalytic C23 residue while preserving OTUB1 deubiquitinase activity. Furthermore, incorporation of this recruiter into bifunctional molecules enabled the development of active CFTR-targeting DUBTACs, compound **25** (MS2134). Together, these studies establish a new OTUB1 recruiter scaffold and provide additional molecular tools for DUBTAC-mediated targeted protein stabilization.

## RESULTS AND DISCUSSION

### Discovery of a Novel Four-Membered Ring OTUB1 Chemotype

The discovery of EN523 established OTUB1 as a tractable recruiter for DUBTAC-mediated targeted protein stabilization.^9^ However, despite growing interest in DUBTAC technology, only a limited number of OTUB1 recruiter chemotypes have been reported to date, leaving the breadth of OTUB1 ligandability largely unexplored. Building upon our previous identification of MS5105 and MS8572,^12, 20^ we sought to further expand the ligandable chemical space of OTUB1 through scaffold diversification. To this end, a series of analogues containing distinct heterocyclic cores were designed and evaluated for their ability to covalently engage OTUB1 using an intact protein mass spectrometry assay.^12^

Based on MS8572, we first explored the middle heterocyclic ring (**Figure 2A**). Initial scaffold exploration revealed a strong dependence of OTUB1 engagement on ring size. Therefore, a series of spirocyclic analogues (compounds **1** – **4**) were synthesized and evaluated. Compound **1**, with a 2,6-diazaspiro[3.3]heptane ring, showed approximately 29% covalent modification of OTUB1. However, increasing either ring size within the spirocyclic system attenuated the modification efficiency (compounds **2** - **4**). The carbonyl group also appeared to play an important role, as its removal resulted in an approximately 3-fold reduction in modification rate (compound **5**: ∼10% modification), consistent with our previous structure-activity relationship (SAR) studies.^20^ In addition to the spirocyclic systems, we designed and synthesized a series of 4-membered-ring analogues, represented by compounds **6** – **9**. Among these, compound **6**, featuring a *p*-amino azetidinyl moiety, showed the highest modification rate (∼36%) (**Figure 2A and B**). Interestingly, reversing the orientation of this moiety, such that the azetidinyl group was directly attached to the carbonyl group, substantially reduced the modification efficiency (compound **7**: ∼ 8%). Furthermore, *N*-methylation of the amide in compound **6** nearly abolished activity (compound **8**, <3% modification). Extending the nitrogen outside the ring also had a detrimental effect, with compound **9** showing < 3% modification.

**Figure 2.**
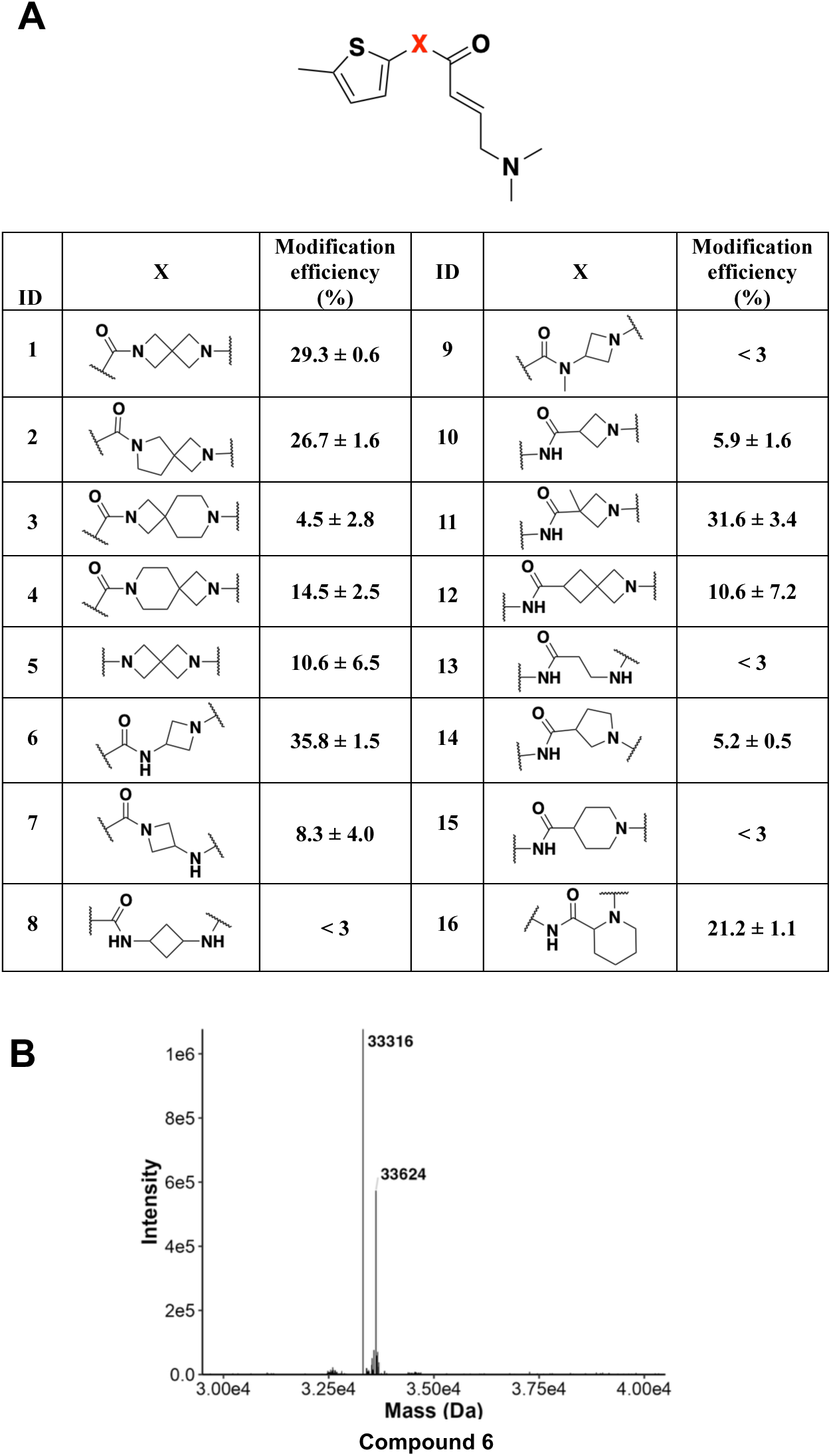
SAR results of the middle heterocyclic ring scaffold. (A) Chemical structures of compounds **1** – **16** and their effects on covalent modification of OTUB1. OTUB1-ligand adduct formation was quantified by intact protein mass spectrometry following incubation of recombinant OTUB1 (10 μM) with each compound at a 250:1 ligand-to-protein molar ratio for 1 h. The percentage of OTUB1 modification was calculated as follows: (%) = (OTUB1-ligand adduct/(OTUB1 + OTUB1-ligand adduct)) × 100. Data are presented as mean ± SD from two independent experiments. (B) Representative mass spectrum of compound **6**.

Since compound **6** represents a novel scaffold with moderate modification efficiency, we next kept the key azetidinyl moiety but reversed the amide orientation, in which the amide nitrogen was linked to the thiophene moiety and the carbonyl group was connected to the azetidinyl moiety. However, the resulting compound **10** showed minimal effect (∼6%). Interestingly, introducing a methyl group at the 3-position of the azetidinyl moiety (**11**) dramatically improved the modification efficiency to ∼32%. Replacing the azetidinyl group with a 2-azaspiro[3.3]heptyl group (compound **12**) provided a modest improvement, with the modification efficiency increasing to ∼11%. Whereas opening the azetidine ring to give the corresponding linear-chain analogue (**13**) resulted in a substantial loss of activity, with modification efficiency dropping below 3%. Next, we investigated the effect of ring size. Both the pyrrolidinyl analogue **14** and the piperidinyl analogue **15** showed reduced potency. Notably, the position of the nitrogen within the ring also influenced modification efficiency, as compound **16**, bearing a piperidine-2-carboxylic acid-derived moiety, achieved ∼21% modification. Collectively, these results indicate that the ring size and orientation of the central heterocycle, together with the presence and positioning of the carbonyl group, are critical determinants of efficient covalent modification of OTUB1.

### Balancing Covalent Reactivity and Selectivity through Warhead Optimization

Having identified compound **6** as a promising lead scaffold, we next explored the impact of covalent warhead modifications on OTUB1 engagement (**Figure 3**). Replacement of the dimethylamino-2-enamide moiety in compound **6** with a dimethylamino-2-alkynamide warhead generated compound **17**, which exhibited a substantial enhancement in OTUB1 modification efficiency, reaching 89.5% (**Figure 3B**). However, this increased reactivity was accompanied by the formation of approximately 30% over-modification, suggesting that the alkynamide warhead may be excessively reactive under the assay conditions. To improve the balance between modification efficiency and labeling selectivity, we next introduced a more sterically demanding dimethylamino-4-methylpent-2-alkynamide warhead, affording compound **18**. Although this modification resulted in a moderate reduction in OTUB1 modification efficiency to 66.5% (**Figure 3B**), it substantially suppressed the formation of over-modified products. These findings indicate that careful tuning of warhead reactivity can significantly improve labeling selectivity while maintaining robust OTUB1 engagement. Consequently, compound **18** was selected as the preferred scaffold for subsequent optimization studies.

**Figure 3.**
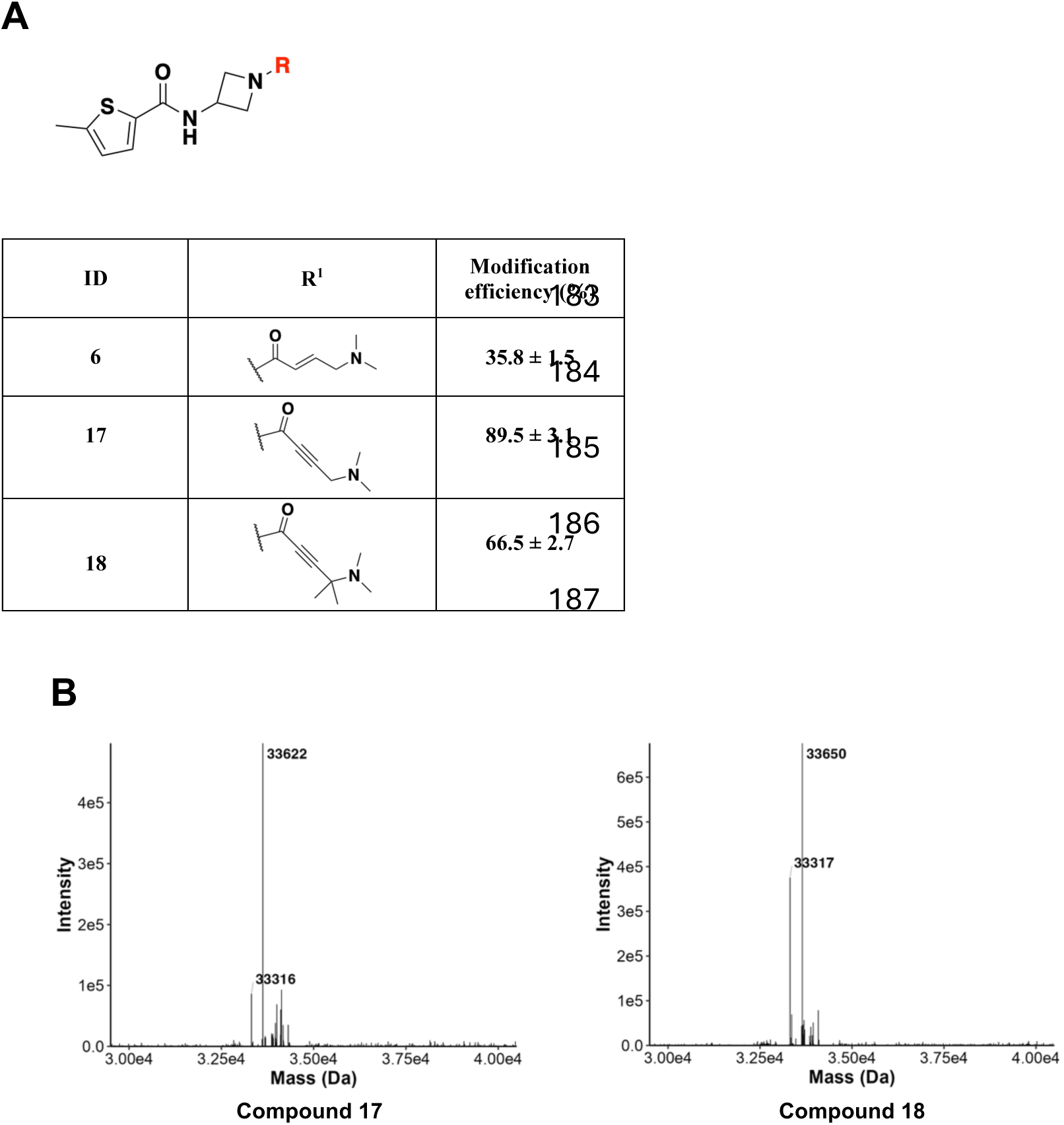
SAR results of the covalent warheads. (A) Chemical structures of compounds **17** and **18** and their effects on covalent modification of OTUB1. OTUB1-ligand adduct formation was quantified by intact protein mass spectrometry following incubation of recombinant OTUB1 (10 μM) with each compound at a 250:1 ligand-to-protein molar ratio for 1 h. The percentage of OTUB1 modification was calculated as follows: (%) = (OTUB1-ligand adduct/(OTUB1 + OTUB1-ligand adduct)) × 100. Data are presented as mean ± SD from two independent experiments. (B) Representative mass spectra of compounds **17** and **18**.

Notably, comparison of compounds **17** and **18** reveals an important design principle for covalent recruiter development: maximal protein labeling does not necessarily translate into an optimal recruiter. Although increasing electrophile reactivity can enhance target engagement, excessive reactivity may also promote nonspecific protein modification and compromise selectivity. Consequently, successful covalent recruiter development requires a careful balance between covalent reactivity and productive molecular recognition, a principle that has been widely recognized in covalent ligand discovery.^22, 23^ Our findings demonstrate that fine-tuning warhead structure can improve labeling selectivity while maintaining robust OTUB1 engagement.

### Optimization of the Left-hand Side (LHS) Aromatic Region

Having identified compound **18** as a suitable scaffold for further optimization, we next investigated the impact of structural modifications within the LHS aromatic region on OTUB1 engagement (**Figure 4**). A series of analogues bearing diverse thiophene-derived substituents were synthesized and evaluated using an intact protein mass spectrometry assay.

**Figure 4.**
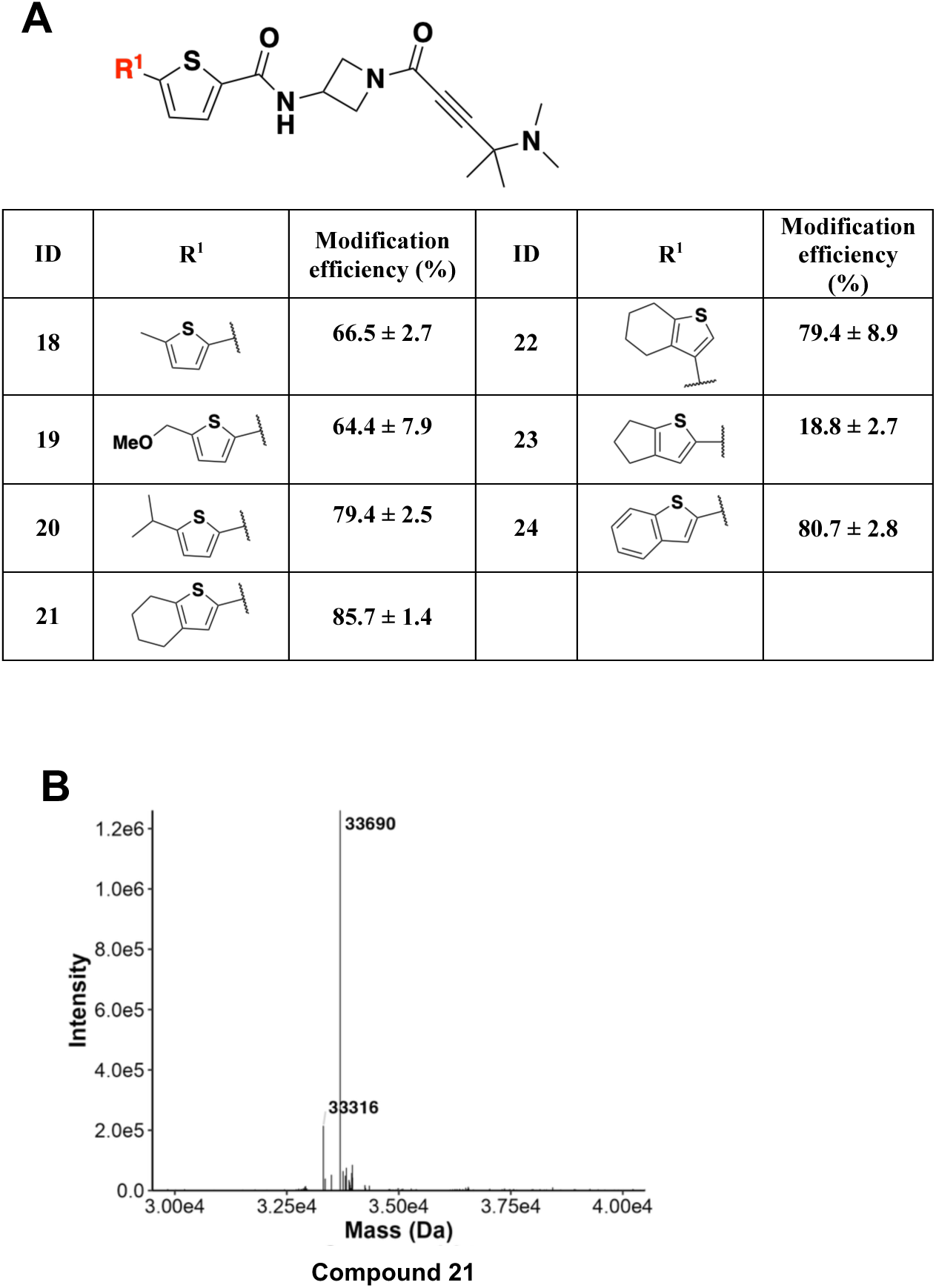
SAR results of the LHS region. (A) Chemical structures of compounds **18** – **24** and their effects on covalent modification of OTUB1. OTUB1-ligand adduct formation was quantified by intact protein mass spectrometry following incubation of recombinant OTUB1 (10 μM) with each compound at a 250:1 ligand-to-protein molar ratio for 1 h. The percentage of OTUB1 modification was calculated as follows: (%) = (OTUB1-ligand adduct/(OTUB1 + OTUB1-ligand adduct)) × 100. Data are presented as mean ± SD from two independent experiments. (B) Representative mass spectrum of compound **21**.

Initial efforts focused on modulating the steric and hydrophobic properties of the thiophene substituent. Introduction of a more polar methoxy substituent (compound **19**) did not affect the OTUB1 modification efficiency. However, replacement of the methyl group with a bulkier isopropyl substituent afforded compound **20**, resulting in an increased OTUB1 modification efficiency of 79.4%. Encouraged by these results, we further explored cyclic substituents to optimize interactions within this region. Compound **21** with a 4,5,6,7-tetrahydrobenzo[*b*]thiophene group exhibited an OTUB1 modification efficiency of 85.7% (**Figure 4B**). Relocation of the carbonyl linkage from the 2-position to the 3-position of the thiophene ring (compound **22**) was tolerated with slightly decreased potency (79.4%). Contraction of the six-membered cyclic substituent in compound **21** to a five-membered analogue (compound **23**) significantly reduced OTUB1 modification efficiency, indicating that substituent ring size contributes to optimal OTUB1 engagement. Lastly, we synthesized the unsaturated benzo[*b*]thiophene scaffold (compound **24**), and tested its modification potency. This compound also maintained high modification efficiency (80.7%). Taken together, these studies identified compound **21** as one of the most potent OTUB1-modifying compounds without observed over-modification. It was selected for further characterization.

### Further Characterization of Compound 21

To further characterize OTUB1 engagement by compound **21**, we first examined its concentration-and time-dependent labeling behavior using EN523 as a positive control. Dose-response studies revealed a progressive increase in OTUB1 modification with increasing compound concentrations, and compound **21** was more efficient than EN523 (**Figure 5A**). Likewise, time-course experiments demonstrated a gradual accumulation of covalent labeling over time (**Figure 5B**), consistent with efficient and sustained engagement of OTUB1. Notably, compound **21** exhibited faster labeling kinetics than EN523, with >50% of OTUB1 modified within 1 h, compared with more than 8 h required for EN523 to achieve a similar level of modification. These findings further validate the enhanced target engagement observed during the SAR campaign and demonstrate that the four-membered ring scaffold can support robust covalent recruitment of OTUB1.

**Figure 5.**
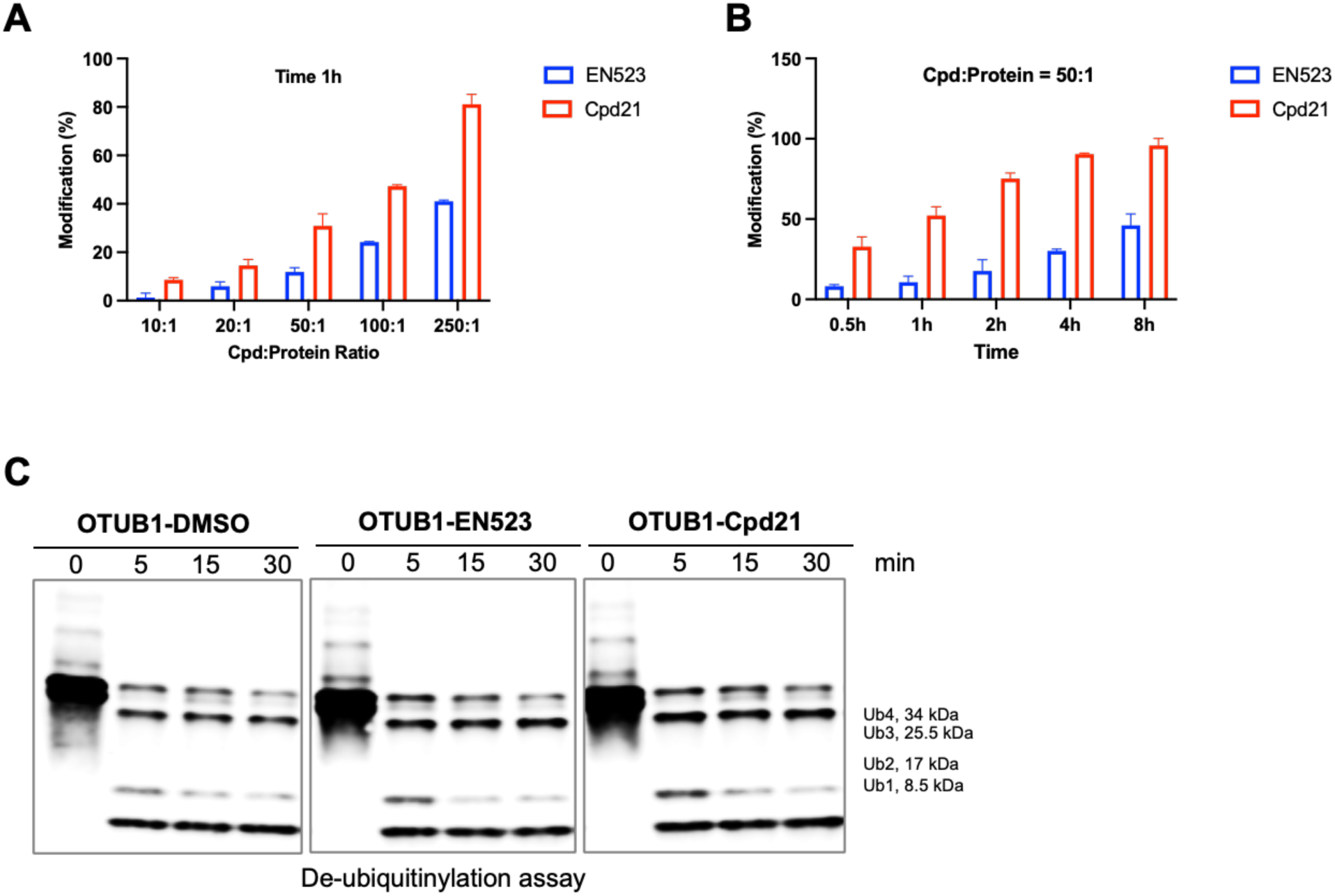
Functional characterization of compound 21. (A) Dose-dependent covalent modification of OTUB1 by compound **21** determined by intact protein mass spectrometry. Recombinant OTUB1 (10 μM) was incubated with compound **21** at the indicated ligand-to-protein molar ratios (10:1–250:1) for 1 h, and OTUB1-ligand adduct formation was quantified by intact protein mass spectrometry. Data are presented as mean ± SD from two independent experiments. (B) Time-dependent covalent modification of OTUB1 by compound **21** determined by intact protein mass spectrometry. Recombinant OTUB1 (10 μM) was incubated with compound **21** at a 50:1 ligand-to-protein molar ratio, and OTUB1 modification was monitored at the indicated time points. Data are presented as mean ± SD from two independent experiments. (C) Evaluation of OTUB1 deubiquitinase activity following covalent modification by compound **21**. Recombinant OTUB1 was preincubated with DMSO, EN523, or compound **21** and subsequently incubated with K48-linked tetra-ubiquitin at 37 °C. Deubiquitination was monitored at the indicated time points by immunoblot analysis.

Next, we sought to determine whether covalent engagement of OTUB1 by compound **21** is compatible with the functional requirements of DUBTAC-mediated protein stabilization. Unlike conventional covalent inhibitors, DUBTAC recruiters must maintain efficient target engagement while preserving the catalytic activity of the recruited deubiquitinase. Previous studies established that the covalent OTUB1 ligand EN523 selectively targets the non-catalytic C23 residue without impairing OTUB1 enzymatic activity, thereby enabling OTUB1-dependent targeted protein stabilization through the DUBTAC platform.^9, 12, 20^ Accordingly, preservation of OTUB1 activity represents a critical criterion for the evaluation of new OTUB1 recruiter chemotypes. As shown in **Figure 5C**, compound **21** did not measurably impair OTUB1 enzymatic function following covalent engagement. This observation is consistent with previous structural and mechanistic studies demonstrating that C23 is located outside the catalytic center of OTUB1 and can therefore be targeted without disrupting deubiquitinase activity.^19^ Importantly, maintenance of OTUB1 function distinguishes productive recruiters from inhibitory covalent ligands and is essential for DUBTAC-mediated protein stabilization, as inhibition of the recruited deubiquitinase would be expected to compromise target deubiquitination and stabilization.

Collectively, these findings demonstrate that compound **21** combines efficient and sustained OTUB1 engagement with preservation of enzymatic activity, thereby establishing it as a functional OTUB1 recruiter suitable for DUBTAC development.

### Selective Engagement of OTUB1 by Compound 21

A key challenge in the optimization of covalent ligands is distinguishing genuine improvements in target engagement from nonspecific increases in electrophilic reactivity.^23^ Given the substantially enhanced OTUB1 labeling efficiency observed for compound **21**, we next sought to determine whether this improvement arose from productive target recognition or simply reflected broader covalent reactivity toward cysteine-containing proteins.

To assess protein selectivity, compound **21** was evaluated against a panel of structurally and functionally distinct cysteine-containing proteins, including full-length OTUB1 (OTUB1 FL), the OTUB1 C23S mutant (OTUB1 C23S Mu), the catalytic domain of G9a-like protein (GLP; residues 982–1266),^24^ and the DNA-binding domain of interferon regulatory factor 3 (IRF3-DBD),^25^ using intact protein mass spectrometry. As shown in **Figure 6**, compound **21** efficiently modified OTUB1 FL, achieving 52.2 ± 5.5% protein labeling under the condition of compound/protein ratio = 50:1 and 1 h incubation time. In contrast, no detectable modification was observed for the OTUB1 C23S mutant, GLP catalytic domain, or IRF3-DBD under identical experimental conditions. The complete loss of labeling upon mutation of C23 confirms that covalent engagement is strictly dependent on the non-catalytic C23 residue, which has previously been established as the ligandable hotspot exploited by OTUB1 recruiters for DUBTAC development.^9, 12, 20^ Importantly, despite the presence of cysteine residues in GLP catalytic domain and IRF3-DBD, no detectable labeling was observed for either protein. These findings indicate that the enhanced OTUB1 engagement achieved through scaffold optimization is not accompanied by increased nonspecific electrophilic reactivity. Rather, the improved labeling efficiency appears to result from productive molecular recognition and favorable interactions between compound **21** and OTUB1.

**Figure 6.**
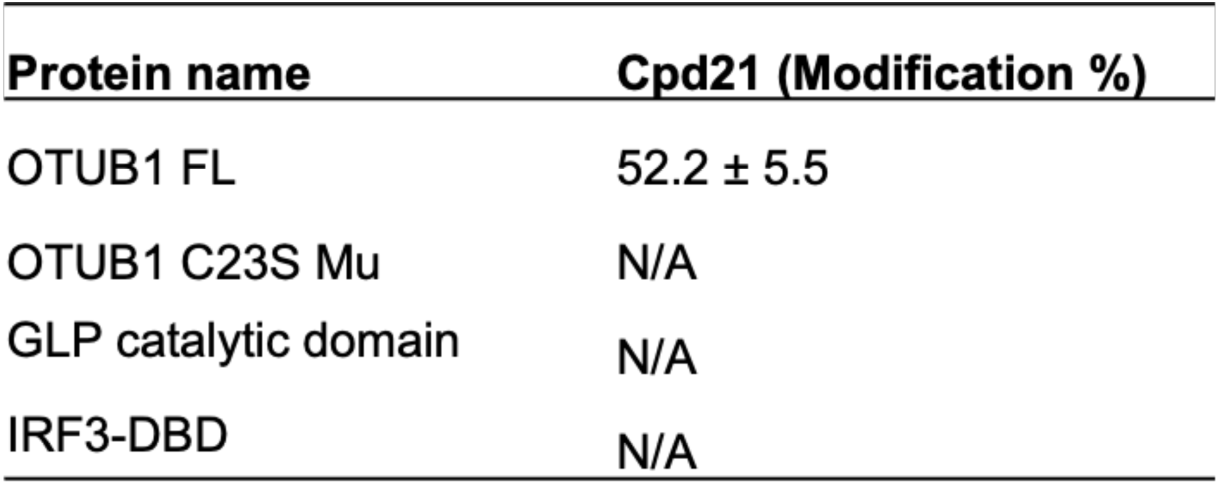
Selective engagement of OTUB1 by compound 21. Covalent modification efficiencies of compound **21** toward recombinant OTUB1 full-length protein (OTUB1 FL), OTUB1 C23S mutant, the catalytic domain of G9a-like protein, and interferon regulatory factor 3 DNA-binding domain (IRF3-DBD) determined by intact protein mass spectrometry. Proteins (10 μM) were incubated with compound **21** at a 50:1 ligand-to-protein molar ratio for 1 h. The percentage of protein-ligand adduct formation was calculated as follows: (%) = (protein-ligand adduct/(protein + protein-ligand adduct)) × 100. Data are presented as mean ± SD from two independent experiments.

### Development of compound 21 based CFTR DUBTACs

Having established compound **21** as an improved OTUB1 recruiter, we next sought to determine whether this newly identified chemotype could be translated into functional DUBTACs capable of mediating targeted protein stabilization (TPS). As the first successful demonstration of DUBTAC technology utilized mutant cystic fibrosis transmembrane conductance regulator (ΔF508-CFTR) as the target substrate,^9^ and because CFTR stabilization remains the most extensively validated OTUB1-dependent DUBTAC model reported to date,^12, 20^ we selected ΔF508-CFTR as a benchmark system to evaluate the utility of compound **21**.

The ΔF508 mutation represents the most common disease-causing CFTR variant in patients with cystic fibrosis and results in protein misfolding, ubiquitination, and rapid proteasomal degradation.^26^ Importantly, the clinical success of FDA-approved CFTR modulators, including lumacaftor, tezacaftor, elexacaftor, and ivacaftor, has demonstrated that restoration of ΔF508-CFTR function can provide substantial therapeutic benefit.^27, 28^ Consequently, restoration of ΔF508-CFTR protein levels through targeted stabilization has emerged as an attractive therapeutic strategy and provides a stringent functional assay for evaluating DUBTAC activity.^9, 12, 20^

To investigate whether the newly developed compound **21** could support DUBTAC-mediated protein stabilization, we introduced a nitrogen atom into the six-membered ring adjacent to the thiophene moiety to provide an exit vector. Next, we linked this compound **21** analogue to the CFTR-binding ligand lumacaftor through an eight-carbon linker to afford compound **25** (MS2134) (**Figure 7A**). This compound was subsequently evaluated in CFBE41o−4.7 human bronchial epithelial cells expressing ΔF508-CFTR. We first tested the impact of compound **25** on CFTR protein levels at multiple concentrations. As shown in **Figure 7B**, compound **25** robustly increased CFTR protein levels at concentrations as low as 2 μM. Next, in a related time-course study, compound **25** started to stabilize CFTR after 12 h of treatment, with the maximal stabilization effect observed at 18 h (**Figure 7C**). Overall, compound **25** induced robust accumulation of ΔF508-CFTR protein, providing direct evidence that the newly identified four-membered-ring OTUB1 recruiter can be successfully incorporated into a functional DUBTAC.

**Figure 7.**
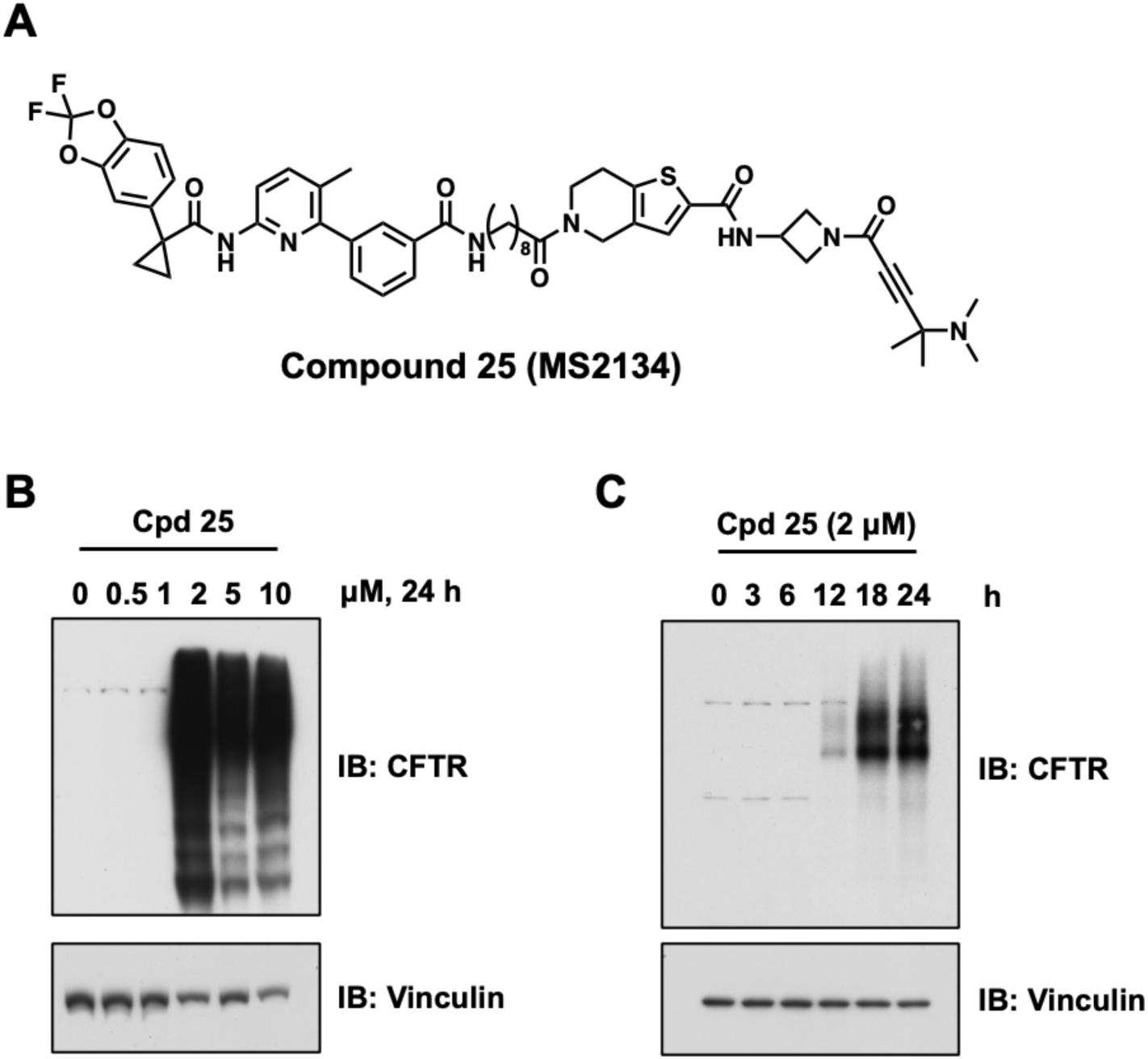
Discovery of compound 25, an OTUB1-based DUBTAC derived from compound 21. (A) Chemical structure of compound 25. (B) WB analysis of compound **25**’s effect on stabilizing the CFTR protein level in ΔF508-CFTR cells treated with compound **25** for 24 h at the indicated concentrations. (C) WB analysis of compound **25**’s effect on stabilizing the CFTR protein level in ΔF508-CFTR cells treated with compound **25** at 2 µM for the indicated times. WB results in panels B and C are representative of two independent experiments. Vinculin was used as a loading control.

## CHEMICAL SYNTHESIS

Compounds **1** – **4,** and **6** – **9** were synthesized following similar procedures for preparing MS8572 (**Scheme 1**).^20^ Starting from commercially available **I-1** and various mono-*N*-Boc-protected diamines, through condensation reaction followed by removal of the Boc protecting group under acid conditions, intermediates **I-2** – **I-9** were obtained. These intermediates were then conjugated with (*E*)-4-(dimethylamino)but-2-enoic acid to yield desired compounds in moderate yields.

**Scheme 1.**
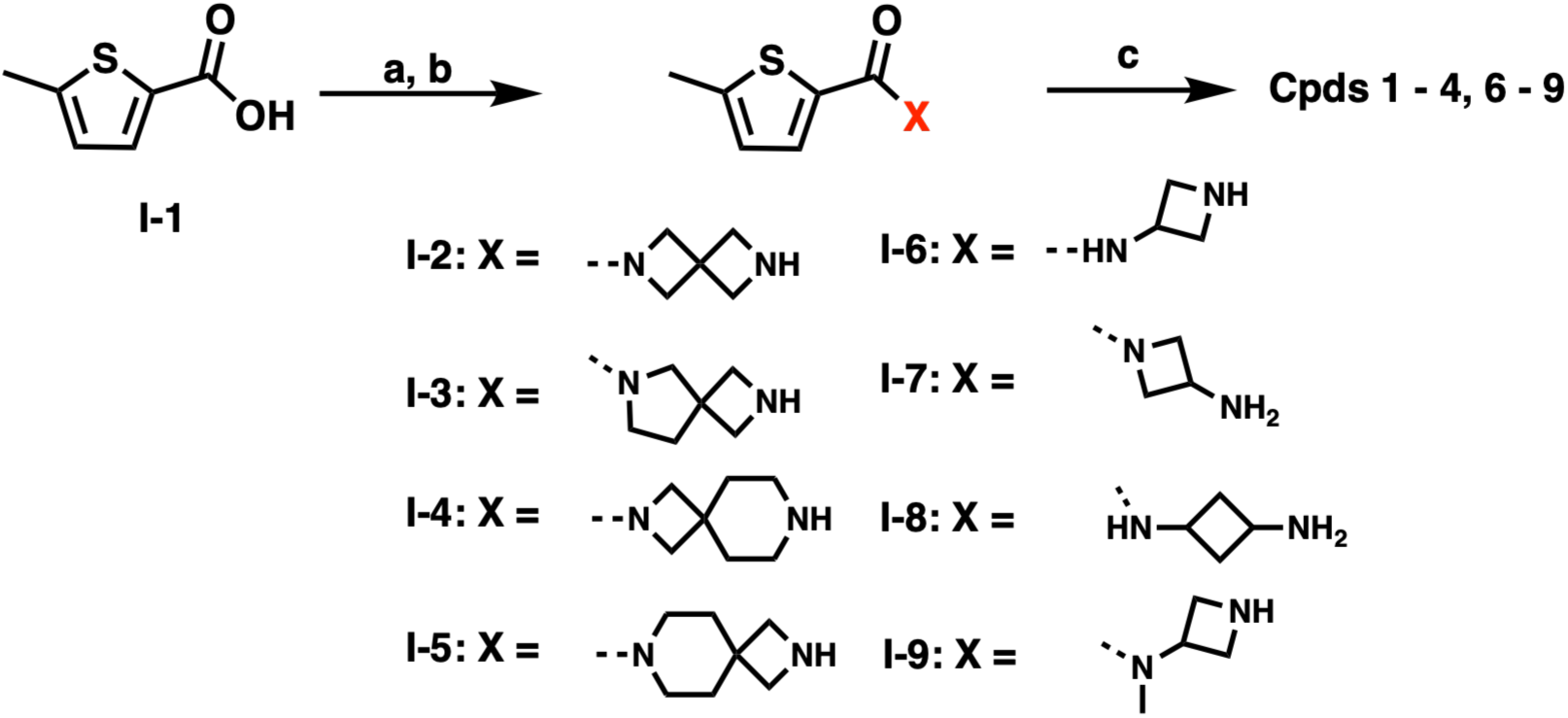
Synthesis of compounds **1** – **4** and **6 - 9***^a^ ^a^*Reation and conditions: (a) various mono-*N*-Boc-protected diamine, HATU, DIEA, DMF, rt; (b) TFA, DCM, rt; (c) (*E*)-4-(dimethylamino)but-2-enoic acid, HATU, DIEA, DMF, rt, 12 - 56% yield over 3 steps.

The synthetic route for compound 5 was illustrated in **scheme 2**. Starting from 2-bromo-5-methylthiophene (**I-10**), Buchwald–Hartwig cross coupling with *tert*-butyl 2,6-diazaspiro[3.3]heptane-2-carboxylate, followed by acid-mediated removal of the Boc protecting group, afforded intermediates **I-11**. Subsequent coupling of **I-11** with (*E*)-4-(dimethylamino)but-2-enoic acid furnished compound **5**.

**Scheme 2.**
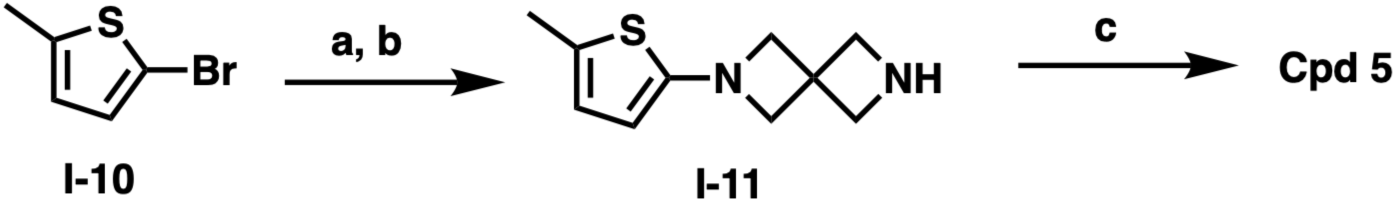
Synthesis of compound **5***^a^ ^a^*Reation and conditions: (a) *tert-*butyl 2,6-diazaspiro[3.3]heptane-2-carboxylate, Pd(PPh_3_)_2_Cl_2_, K_3_PO_4_, 1,4-dioxane, H_2_O 100°C; (b) TFA, DCM, rt; (c) (*E*)-4-(dimethylamino)but-2-enoic acid, HATU, DIEA, DMF, rt, 23% yield over 3 steps;

**Scheme 3.**
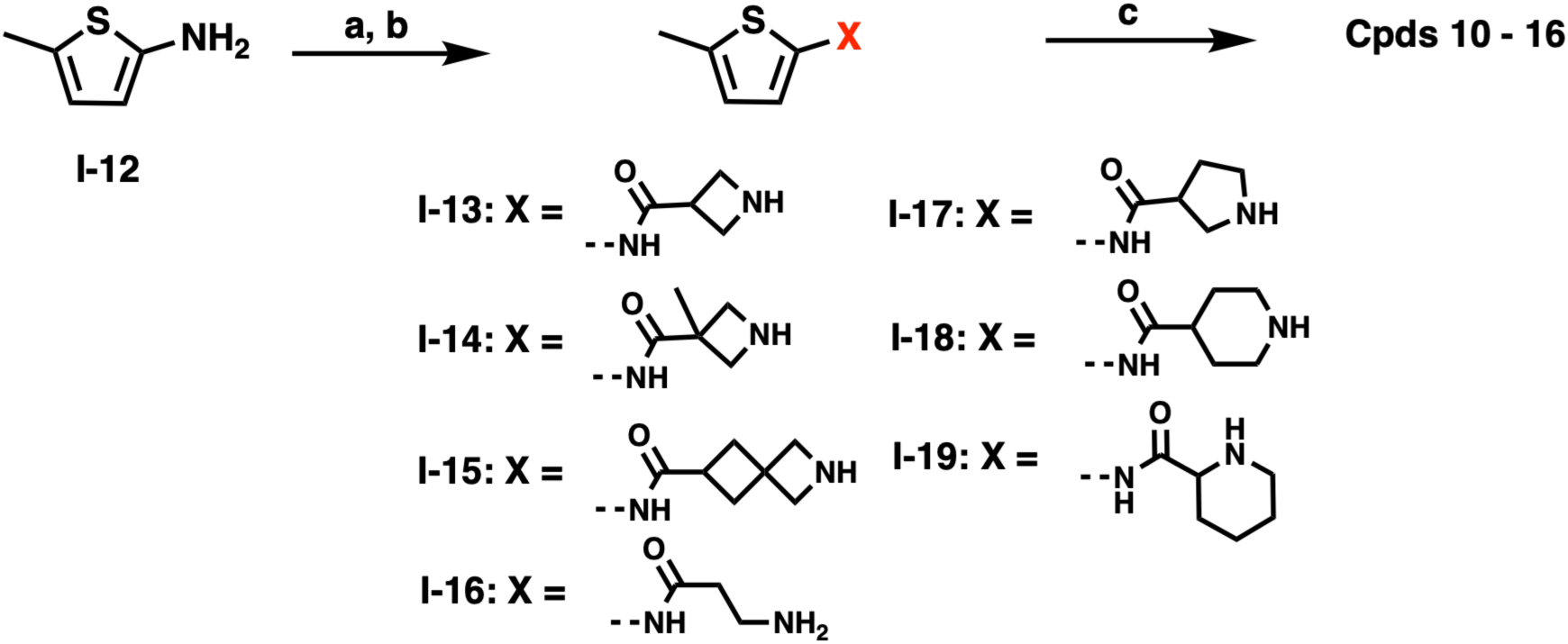
Synthesis of compounds **10** – **16***^a^ ^a^*Reation and conditions: (a) various *N*-Boc-protected amino acid, HATU, DIEA, DMF, rt; (b) TFA, DCM, rt, 57 - 85% yield over two-steps; (c) (*E*)-4-(dimethylamino)but-2-enoic acid, HATU, DIEA, DMF, rt, 16 - 43% yield.

Similarly, starting from commercially available intermediates **I-10**, intermediates **I-13** – **I-19** were obtained in moderate to good yields through condensation reaction followed by removal of the Boc protecting group under acid condition. These intermediates were then conjugated with (*E*)-4-(dimethylamino)but-2-enoic acid to yield compounds **10** – **16** (**Scheme 3**).

The synthetic routes for compounds **17 and 18** are showed in **Scheme 4**. Intermediate **I-10** was subjected to a condensation reaction with 4-(dimethylamino)but-2-ynoic acid or 4-(dimethylamino)-4-methylpent-2-ynoic acid to obtain compounds **17** or **18**.

**Scheme 4.**
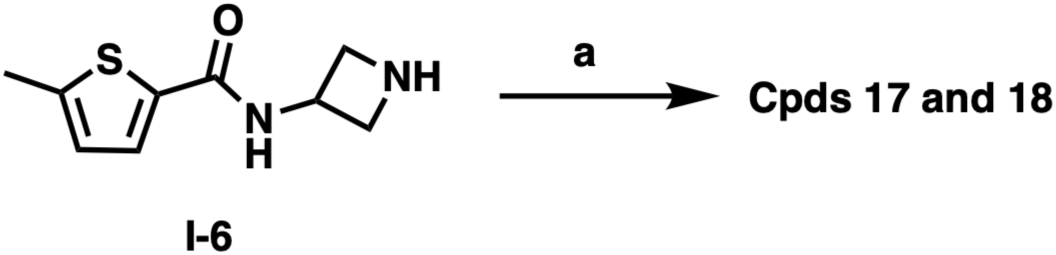
Synthesis of compounds **17** and **18***^a^ ^a^*Reation and conditions: (a) 4-(dimethylamino)but-2-ynoic acid or 4-(dimethylamino)-4- methylpent-2-ynoic acid, HATU, DIEA, DMF, rt, 28 - 31% yield.

**Scheme 5.**
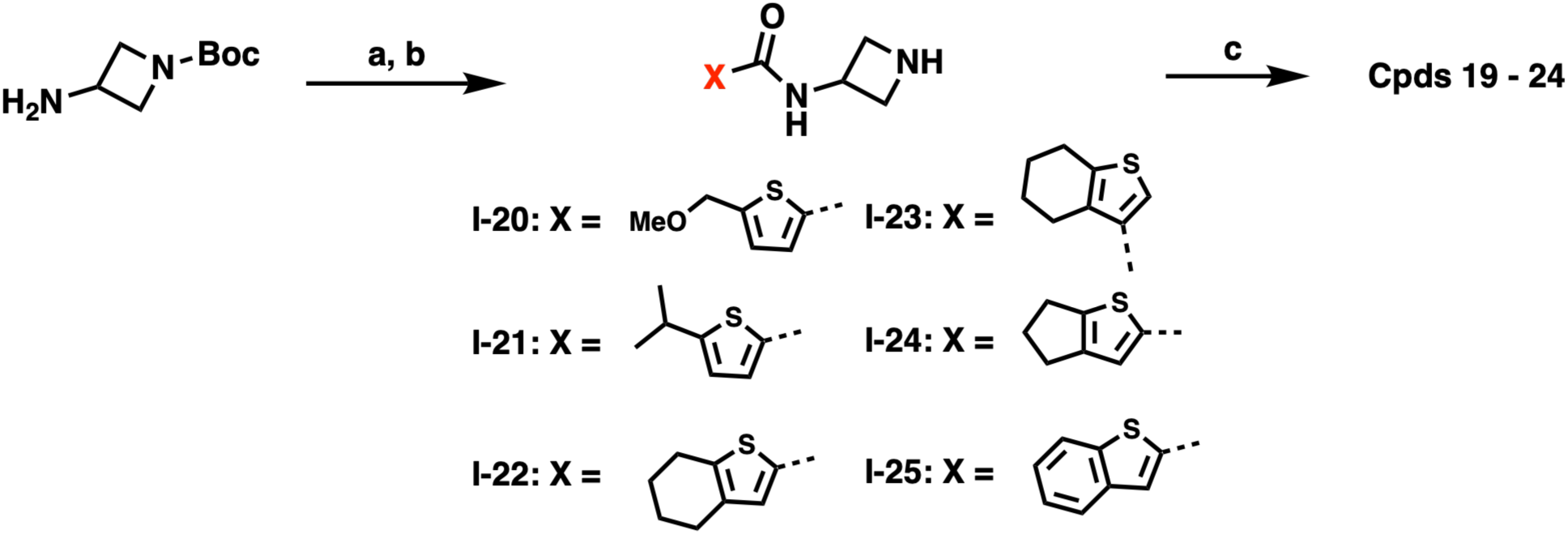
Synthesis of compounds **19** – **24***^a^ ^a^*Reation and conditions: (a) various thiophene carboxylic acids, HATU, DIEA, DMF, rt; (b) TFA, DCM, rt, 37 - 63% yield over two-steps; (c) 4-(dimethylamino)-4-methylpent-2-ynoic acid, HATU, DIEA, DMF, rt, 19 - 41%.

The synthetic routes for compounds **19** – **24** are illustrated in **Scheme 5**. Starting from commercially available intermediate **I-24**, conjugation with various thiophene carboxylic acids followed by TFA-mediated deprotection and subsequent installation of 4-(dimethylamino)-4-methylpent-2-ynoic acid in the final step afforded the desired compounds.

The synthetic route for compound **25** is illustrated in **Scheme 6**. Starting from commercially available intermediate **I-26**, coupling with benzyl 3-aminoazetidine-1-carboxylate, followed by Pd/C-mediated deprotection afforded **I-27**. **I-27** was further converted to **I-28** by reacting with 4-(dimethylamino)-4-methylpent-2-ynoic acid. **I-28** was then reacted with the previously reported intermediate **I-29**^11^ via a regular coupling reaction to get the final DUBTACs **25**.

**Scheme 6.**
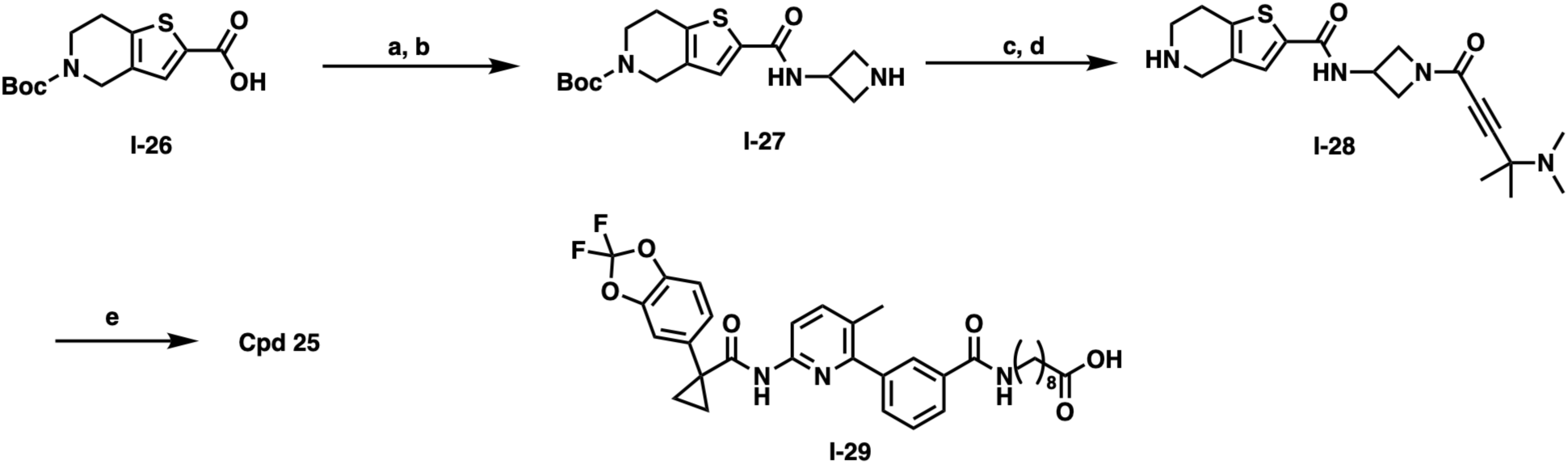
Synthesis of compounds **25***^a^ ^a^*Reation and conditions: (a) benzyl 3-aminoazetidine-1-carboxylate, HATU, DIEA, DMF, rt; (b) Pd/C, triethylsilane, 50°C, MeOH, 15% yield over 2-steps; (c) 4-(dimethylamino)-4-methylpent-2-ynoic acid, HATU, DIEA, DMF, rt; (d) TFA, DCM, rt, 56% yield over 2-steps; (e) **I-29**, HATU, DIEA, DMF, rt, 24% yield.

## CONCLUSIONS

In summary, we report the discovery and optimization of a novel four-membered ring OTUB1 chemotype as a new class of covalent OTUB1 recruiters. Through systematic SAR studies, we identified compound **21** as a highly efficient ligand that selectively engages the non-catalytic C23 residue in OTUB1. Biochemical characterization further demonstrated that compound **21** exhibits robust concentration-and time-dependent OTUB1 engagement while preserving OTUB1 deubiquitinase activity, a key requirement for DUBTAC-mediated targeted protein stabilization. Importantly, incorporation of this recruiter scaffold into bifunctional molecules enabled the development of functional OTUB1-based DUBTACs capable of stabilizing ΔF508-CFTR in cells, demonstrating the utility of this scaffold for targeted protein stabilization.

Together with previously reported OTUB1 recruiters, the four-membered ring scaffold described here further expands the repertoire of ligands available for OTUB1-directed targeted protein stabilization. The successful stabilization of ΔF508-CFTR demonstrates the utility of this scaffold for DUBTAC development and further supports OTUB1 as a versatile platform for targeted protein stabilization. Expansion of OTUB1 recruiter chemotypes may facilitate the development of future DUBTACs with improved potency, selectivity, and pharmacological properties.

## EXPERIMENTAL SECTION

### Expression and Purification of Recombinant Proteins

Human OTUB1 wild-type (NM_017670.3) and C23S mutant were cloned, expressed and purified according to a previously described protocol.^12^ The catalytic domain of human GLP (amino acid residues 982–1266) was cloned, expressed, and purified according to a previously described protocol.^24^

The DNA-binding domain of human interferon regulatory factor 3 (IRF3-DBD) was cloned into the pET15b expression vector containing an N-terminal His tag, expressed in BL21-CodonPlus (DE3)-RIPL cells (Agilent Technologies), and purified by Ni-NTA affinity chromatography followed by size-exclusion chromatography. Purified protein was concentrated, aliquoted, flash-frozen in liquid nitrogen, and stored at −80 °C until use. The amino acid sequence of the recombinant IRF3-DBD construct is provided below: MGTPKPRILPWLVSQLDLGQLEGVAWVNKSRTRFRIPWKHGLRQDAQQEDFGIFQAWAEATGAYVPGCDKPDLPTWKRNFRSALNRKEGLRLAEDRSKDPHDPHKIYEFVNSG

### Intact Protein Mass Spectrometry-Based SAR Analysis

Recombinant OTUB1 protein was incubated with the indicated compounds in reaction buffer containing 50 mM Tris-HCl (pH 7.5), 150 mM NaCl, and 0.5 mM TCEP. Unless otherwise indicated, reactions were carried out using 10 μM OTUB1 and a 250:1 ligand-to-protein molar ratio for 1 h at room temperature. Following incubation, samples were buffer exchanged into 50 mM ammonium bicarbonate using a 10 kDa molecular weight cutoff (MWCO) centrifugal filter unit (Millipore) and diluted in MS buffer containing 30% acetonitrile and 0.2% formic acid to a final protein concentration of 10 μM.

Intact protein mass spectrometric analysis was performed using either an Agilent LC/MSD Time-of-Flight (TOF) mass spectrometer (Agilent Technologies) equipped with an electrospray ionization source or an Acquity H-Class UPLC coupled to a Xevo G2 XS QTOF mass spectrometer (Waters) operated in positive ion mode. Protein masses were deconvoluted and analyzed using TOF Protein Confirmation Software (Agilent BioConfirm version 2.0 or 11.0).

The percentage of protein–ligand adduct formation was calculated according to the following equation: % Adduct Formation = [Protein–Ligand Adduct / (Unmodified Protein + Protein–Ligand Adduct)] × 100

For SAR studies, data are presented as mean ± SD from two independent experiments.

### Selectivity Assessment by Intact Protein Mass Spectrometry

To evaluate selectivity, compound **21** was incubated with recombinant full-length OTUB1, OTUB1 C23S mutant, GLP catalytic domain, and IRF3-DBD under the same assay conditions described above. Each protein (10 μM) was incubated with compound **21** at a 50:1 ligand-to-protein molar ratio for 1 h at room temperature. Protein–ligand adduct formation was assessed by intact protein mass spectrometric analysis as described above. Data are presented as mean ± SD from two independent experiments.

### Concentration-and Time-Dependent OTUB1 Modification Assays

For concentration-dependent OTUB1 modification studies, recombinant OTUB1 protein (10 μM) was incubated with compound **21** at ligand-to-protein molar ratios of 10:1, 20:1, 50:1, 100:1, and 250:1 in reaction buffer for 1 h at room temperature.

For time-course studies, recombinant OTUB1 protein (10 μM) was incubated with compound **21** at a 50:1 ligand-to-protein molar ratio in reaction buffer, and aliquots were collected after 0.5, 1, 2, 4, and 8 h of incubation at room temperature.

Samples were analyzed by intact protein mass spectrometric analysis as described above. OTUB1 modification percentages were calculated based on the relative abundance of unmodified OTUB1 and OTUB1–ligand adducts. Data are presented as mean ± SD from two independent experiments.

### *In Vitro* OTUB1 Deubiquitinase Activity Assay

To assess whether covalent ligand engagement affects OTUB1 enzymatic activity, recombinant OTUB1 protein (10 μM) was preincubated with DMSO, EN523, or compound **21** at a 500:1 ligand-to-protein molar ratio in reaction buffer for 1 h at room temperature. Excess ligand was subsequently removed by buffer exchange using a 10 kDa MWCO centrifugal filter unit (Millipore).

For deubiquitination assays, OTUB1 (1.5 μM) pretreated with DMSO, EN523, or compound **21** was incubated with K48-linked tetra-ubiquitin (3.6 μM, LifeSensors) in reaction buffer at 37 °C for the indicated time points. Reactions were terminated by addition of 4× Laemmli sample buffer containing reducing agent followed by heating at 95 °C for 6 min.

Reaction products were separated by SDS-PAGE and analyzed by immunoblotting using an anti-ubiquitin antibody (Ubiquitin (P4D1) Mouse mAb, #3936S, Cell Signaling Technology). An IRDye 680RD donkey anti-mouse IgG secondary antibody (#926-68072, LI-COR) was used for signal detection. Protein signals were visualized using an Odyssey CLx imaging system (LI-COR). OTUB1 deubiquitinase activity was evaluated by monitoring the disappearance of K48-linked polyubiquitin substrate and the generation of lower-molecular-weight ubiquitin species.

### Cell Culture

CFBE41o−4.7 ΔF508-CFTR human cystic fibrosis bronchial epithelial cells (Sigma, SCC159) were maintained in α-MEM medium (Sigma, M2279) supplemented with 10% fetal bovine serum, 2 mM L-glutamine, 300 μg/mL hygromycin B, and penicillin–streptomycin.^29^ The cell line was generated from parental CFBE41o− cells through stable expression of the ΔF508-CFTR mutant. For compound treatment experiments, cells were seeded in 6-well plates coated with an extracellular matrix mixture consisting of fibronectin (10 μg/mL; Sigma, F2006), PureCol collagen (30 μg/mL; Sigma, 5006), and bovine serum albumin (100 μg/mL; Sigma, 126575). Cells were treated with the indicated compounds after reaching approximately 60–70% confluency.

### Western Blot Analysis of DUBTAC-Mediated ΔF508-CFTR Stabilization

Cells were lysed in EBC (50 mM Tris pH 7.5, 120 mM NaCl, 0.5% NP-40) supplemented with protease inhibitors (Pierce) and phosphatase inhibitors (phosphatase inhibitor cocktail set I and II, Calbiochem), and protein concentrations were measured as described previously ^30^. The lysates (60 µg protein) were resolved by 6% SDS-PAGE at 130 V for 120 min for CFTR or 10% SDS-PAGE at 130 V for 80 min for other proteins, transferred to PVDF membrane, and immunoblotted with the indicated antibodies at 4°C overnight. The membrane was washed 4 times with Tris-buffered saline containing 0.1% Tween-20 (TBST), incubated with secondary antibody for 1 hour at room temperature, and washed 4 times with TBST buffer. The CFTR (#78335, 1:1,000) antibody was obtained from Cell Signaling Technology. The Vinculin antibody (V-4505, 1:50,000), peroxidase-conjugated anti-mouse secondary antibody (A-4416, 1:3000) and peroxidase-conjugated anti-rabbit secondary antibody (A-4914, 1:3,000) were purchased from Sigma.

### Statistical Analysis

Data are presented as mean ± SD from two independent experiments unless otherwise indicated. Statistical analysis was performed using GraphPad Prism. Details of statistical tests are provided in the corresponding figure legends.

### Chemistry General Procedures

All commercial chemical reagents and solvents were used for the reactions without further purification. Flash column chromatography was performed on Teledyne ISCO CombiFlash Rf+ instrument equipped with a 220/254/280 nm wavelength UV detector and a fraction collector. Normal phase column chromatography was conducted on silica gel columns with either hexane/ethyl acetate or dichloromethane/methanol as eluent. Reverse phase column chromatography was conducted on HP C18 RediSep Rf columns, and the gradient was set to 10% of acetonitrile in H_2_O containing 0.1% TFA progressing to 100% of acetonitrile. All final compounds were purified with preparative high-performance liquid chromatography (HPLC) on an Agilent Prep 1290 infinity II series with the UV detector set to 220/254/280 nm at a flow rate of 40 mL/min. Samples were injected onto a Phenomenex Luna 750 x 30 mm, 5 μm C18 column, and the gradient was set to 10% of acetonitrile in H_2_O containing 0.1% TFA progressing to 100% of acetonitrile. LCMS was performed by an Agilent 1200 series system with DAD detector and a 2.1 mm x 150 mm Zorbax 300SB-C18 5 μm column for chromatography and high-resolution mass spectra (HRMS) that were acquired in positive ion mode using an Agilent G6230BA Accurate Mass TOF with an electrospray ionization (ESI) source. Samples (0.8 μL) were injected onto a C18 column at room temperature, and the flow rate was set to 0.6 mL/min with water containing 0.1% formic acid as solvent A and acetonitrile containing 0.1% formic acid as solvent B. Nuclear magnetic resonance (NMR) spectra were acquired on Bruker DRX 400 MHz for proton (^1^H NMR) and 101 MHz for carbon (^13^C NMR). Chemical shifts for all compounds are reported in parts per million (ppm, δ). The format of chemical shift was reported as follows: chemical shift, multiplicity (s = singlet, d = doublet, t = triplet, q = quartet, m = multiplet), coupling constant (J values in Hz), and integration. All final compounds had > 95% purity using the HPLC methods described above.

### (*E*)-4-(dimethylamino)-1-(6-(5-methylthiophene-2-carbonyl)-2,6-diazaspiro[3.3]heptan-2-yl)but-2-en-1-one (1)

To a solution of **I-1** (14.2 mg, 0.1 mmol, 1.0 eq) and *tert*-butyl 2,6-diazaspiro[3.3]heptane-2-carboxylate (19.8 mg, 0.1 mmol, 1.0 eq) in DMF, HATU(41.9 mg, 0.11 mmol, 1.1 eq) and DIEA(0.04 mL, 0.22 mmol, 2.0 eq) were added. The reaction mixture stirred at rt for 60 mins. The resulting crude mixtures were purified via ISCO to yield intermediate. Then the intermediate was dissolved in DCM/TFA (1:1, 1 mL). The reaction mixture was stirred at rt 1 h. After excess TFA was removed, resulting crude mixtures were purified via ISCO to yield intermediate **I-2**. To a solution of **I-2** (6.6 mg, 0.03 mmol, 1.0 eq) and (*E*)-4-(dimethylamino)but-2-enoic acid (3.9 mg, 0.03 mmol, 1.0 eq) in DMF, HATU (12.5 mg, 0.033 mmol, 1.1 eq) and DIEA(13 μL, 0.066 mmol, 2.0 eq) were added. The reaction mixture stirred at rt for 60 mins. The resulting crude mixtures were purified via prep-HPLC to yield titled compound as white solid, 19% yield. ^1^H NMR (400 MHz, Methanol-*d*_4_) δ 7.27 (d, *J* = 4.1 Hz, 1H), 6.75 (d, *J* = 4.6 Hz, 1H), 6.73 – 6.54 (m, 1H), 6.49 – 6.33 (m, 1H), 4.59 (s, 2H), 4.50 – 4.39 (m, 2H), 4.32 – 4.12 (m, 4H), 3.85 (t, *J* = 5.4 Hz, 2H), 2.81 (s, 6H), 2.42 (s, 3H). HRMS (ESI-TOF) m/z: [M+H]^+^ calcd for C_18_H_26_N_3_O_2_S, 348.1740; found: 348.1759.

Compounds **2** - **4** were synthesized following a similar procedure for preparing compound **1**.

### (*E*)-4 (dimethylamino)-1-(6-(5-methylthiophene-2-carbonyl)-2,6-diazaspiro[3.4]octan-2-yl)but-2-en-1-one (2)

White solid, 12% yield. ^1^H NMR (400 MHz, Methanol-*d*_4_) δ 7.40 (d, *J* = 3.8 Hz, 1H), 6.85 – 6.76 (m, 1H), 6.76 – 6.59 (m, 1H), 6.56 – 6.31 (m, 1H), 4.39 – 4.15 (m, 2H), 4.13 – 3.95 (m, 3H), 3.88 (t, *J* = 10.4 Hz, 3H), 3.69 (d, *J* = 53.8 Hz, 2H), 2.85 (s, 6H), 2.51 – 2.43 (m, 3H), 2.32 – 2.08 (m, 2H). HRMS (ESI-TOF) m/z: [M+H]^+^ calcd for C_18_H_26_N_3_O_2_S, 348.1740; found: 348.1763.

### (*E*)-4-(dimethylamino)-1-(2-(5-methylthiophene-2-carbonyl)-2,7-diazaspiro[3.5]nonan-7-yl)but-2-en-1-one (3)

White solid, 31% yield. ^1^H NMR (400 MHz, Methanol-*d*_4_) δ 7.35 – 7.27 (m, 1H), 6.96 – 6.85 (m, 1H), 6.77 (d, *J* = 4.5 Hz, 1H), 6.67 – 6.49 (m, 1H), 4.20 (s, 2H), 3.95 – 3.73 (m, 4H), 3.71 – 3.47 (m, 4H), 2.82 (s, 6H), 2.43 (s, 3H), 1.88 – 1.61 (m, 4H). HRMS (ESI-TOF) m/z: [M+H]^+^ calcd for C_19_H_28_N_3_O_2_S, 362.1897; found: 362.1915.

### (*E*)-4-(dimethylamino)-1-(7-(5-methylthiophene-2-carbonyl)-2,7-diazaspiro[3.5]nonan-2-yl)but-2-en-1-one (4)

White solid, 16% yield. ^1^H NMR (400 MHz, Methanol-*d*_4_) δ 7.15 (d, *J* = 3.7 Hz, 1H), 6.78 – 6.73 (m, 1H), 6.73 – 6.63 (m, 1H), 6.46 (d, *J* = 15.3 Hz, 1H), 4.06 (s, 2H), 3.96 – 3.87 (m, 2H), 3.81 (s, 2H), 3.74 – 3.58 (m, 4H), 2.85 (s, 6H), 2.50 – 2.42 (m, 3H), 1.89 – 1.71 (m, 4H).). HRMS (ESI-TOF) m/z: [M+H]^+^ calcd for C_19_H_28_N_3_O_2_S, 362.1897; found: 362.1910.

### (*E*)-4-(dimethylamino)-1-(6-(5-methylthiophen-2-yl)-2,6-diazaspiro[3.3]heptan-2-yl)but-2-en-1-one (5)

To a solution of the 2-bromo-5-methylthiophene (**I-10**) (177.1 mg, 1 mmol, 1 eq) dissolved in dioxane/H_2_O (4:1, 3 mL), K_3_PO_4_ (424.6 mg, 2 mmol, 2.0 eq), Pd (PPh_3_)_2_Cl_2_ (70.2 mg, 0.1 mmol, 0.1 eq), followed by *tert*-butyl 2,6-diazaspiro[3.3]heptane-2-carboxylate (396.4 mg, 2.0 mmol, 2.0 eq). The reaction mixture was stirred at 100 °C under nitrogen atmosphere overnight. After cooling down to rt, resulting crude mixtures were purified via ISCO to yield intermediate. To a solution of intermediate in DCM (1 mL), TFA (1 mL) was added. The reaction mixture was stirred at rt for 1 h, resulting crude mixtures were purified via ISCO to yield intermediate **I-11**. To a solution of the **I-11** (5.82 mg, 0.03 mmol, 1.0 eq) in DMF (1 mL), HATU (12.5 mg, 0.033 mmol, 1.1 eq), DIEA (35 μL, 0.2 mmol, 2.0 eq) were added followed by (*E*)-4-(dimethylamino)but-2-enoic acid (3.9 mg, 0.03 mmol, 1.0 eq). After stirred at rt for 30 mins, resulting mixtures were purified via prep-HPLC to yield titled compound as white solid, 23% yield. ^1^H NMR (400 MHz, Methanol-*d*_4_) δ 6.69 (s, 2H), 6.55 – 6.32 (m, 2H), 4.50 (s, 2H), 4.22 (d, *J* = 3.2 Hz, 2H), 4.01 – 3.91 (m, 6H), 2.90 – 2.88 (m, 6H), 2.32 (d, *J* = 3.4 Hz, 3H). HRMS (ESI-TOF) m/z: [M+H]^+^ calcd for C_17_H_24_N_3_O_2_S, 334.1584; found: 334.1584.

Compound **6** - **9** were synthesized following similar procedure for preparing compound **1**.

### (*E*)-*N*-(1-(4-(dimethylamino)but-2-enoyl)azetidin-3-yl)-5-methylthiophene-2-carboxamide (6)

White solid, 56% yield. ^1^H NMR (400 MHz, Methanol-*d*_4_) δ 7.47 (d, *J* = 3.7 Hz, 1H), 6.74 (d, *J* = 3.0 Hz, 1H), 6.71 – 6.57 (m, 1H), 6.44 (d, *J* = 15.3 Hz, 1H), 4.76 – 4.63 (m, 1H), 4.58 (t, *J* = 8.6 Hz, 1H), 4.36 – 4.28 (m, 1H), 4.27 – 4.18 (m, 1H), 4.07 – 3.96 (m, 1H), 3.87 (d, *J* = 7.1 Hz, 2H), 2.83 (s, 6H), 2.43 (s, 3H). HRMS (ESI-TOF) m/z: [M+H]^+^ calcd for C_15_H_22_N_3_O_2_S, 308.1427; found: 308.1453.

### (*E*)-4-(dimethylamino)-*N*-(1-(5-methylthiophene-2-carbonyl)azetidin-3-yl)but-2-enamide (7)

White solid, 33% yield. ^1^H NMR (400 MHz, Methanol-*d*_4_) δ 7.29 (d, *J* = 3.1 Hz, 1H), 6.78 (s, 1H), 6.74 – 6.59 (m, 1H), 6.31 (d, *J* = 15.3 Hz, 1H), 4.82 – 4.61 (m, 2H), 4.35 (d, *J* = 38.0 Hz, 2H), 3.98 (s, 1H), 3.88 (d, *J* = 7.2 Hz, 2H), 2.83 (s, 6H), 2.45 (s, 3H). HRMS (ESI-TOF) m/z: [M+H]^+^ calcd for C_15_H_22_N_3_O_2_S, 308.1427; found: 308.1448.

### (*E*)-*N*-(3-(4-(dimethylamino)but-2-enamido)cyclobutyl)-5-methylthiophene-2-carboxamide (8)

White solid, 45% yield. ^1^H NMR (400 MHz, Methanol-*d*_4_) δ 7.48 – 7.35 (m, 1H), 6.71 (s, 1H), 6.69 – 6.54 (m, 1H), 6.34 – 6.20 (m, 1H), 4.46 (p, *J* = 6.9, 6.4 Hz, 1H), 4.38 – 4.26 (m, 1H), 3.89 – 3.77 (m, 2H), 2.80 (s, 6H), 2.73 – 2.61 (m, 1H), 2.47 – 2.35 (m, 4H), 2.35 – 2.25 (m, 1H), 2.08 – 1.91 (m, 1H HRMS (ESI-TOF) m/z: [M+H]^+^ calcd for C_15_H_22_N_3_O_2_S, 322.1584; found: 322.1607.

### *E*)-*N*-(1-(4-(dimethylamino)but-2-enoyl)azetidin-3-yl)-*N*,5-dimethylthiophene-2-carboxamide (9)

White solid, 23% yield. ^1^H NMR (400 MHz, Methanol-*d*_4_) δ 7.54 (d, *J* = 3.7 Hz, 1H), 6.89 – 6.77 (m, 2H), 6.57 (d, *J* = 15.2 Hz, 1H), 4.81 – 4.70 (m, 1H), 4.67 (t, *J* = 8.5 Hz, 1H), 4.45 – 4.35 (m, 1H), 4.34 – 4.24 (m, 1H), 4.14 (d, *J* = 7.2 Hz, 3H), 3.16 (s, 9H), 2.51 (s, 3H). HRMS (ESI-TOF) m/z: [M+H]^+^ calcd for C_16_H_24_N_3_O_2_S, 322.1584; found: 322.1609.

### (*E*)-1-(4-(dimethylamino)but-2-enoyl)-*N*-(5-methylthiophen-2-yl)azetidine-3-carboxamide (10)

To a solution of **I-12** (11.3 mg, 0.1 mmol, 1.0 eq) and 1-(*tert*-butoxycarbonyl)azetidine-3-carboxylic acid (20.1 mg, 0.1 mmol, 1.0 eq) in DMF, HATU (41.9 mg, 0.11 mmol, 1.1 eq) and DIEA (0.04 mL, 0.22 mmol, 2.0 eq) were added. The reaction mixture stirred at rt for 60 mins. The resulting crude mixtures were purified via ISCO to yield intermediate. Then the intermediate was dissolved in DCM/TFA (1:1, 1 mL). The reaction mixture was stirred at rt 1 h. After excess TFA was removed, resulting crude mixtures were purified via ISCO to yield intermediate **I-13**. To a solution of **I-13** (5.9 mg, 0.03 mmol, 1.0 eq) and (*E*)-4-(dimethylamino)but-2-enoic acid (3.9 mg, 0.03 mmol, 1.0 eq) in DMF, HATU(41.9 mg, 0.033 mmol, 1.1 eq) and DIEA(13 μL, 0.066 mmol, 2.0 eq) were added. The reaction mixture stirred at rt for 60 mins. The resulting crude mixtures were purified via prep-HPLC to yield titled compound as white solid, 24% yield. ^1^H NMR (400 MHz, Methanol-*d*_4_) δ 6.80 – 6.62 (m, 1H), 6.53 – 6.37 (m, 3H), 4.54 – 4.39 (m, 2H), 4.29 – 4.20 (m, 1H), 4.19 – 4.11 (m, 1H), 3.95 – 3.85 (m, 2H), 3.63 – 3.51 (m, 1H), 2.87 (s, 6H), 2.35 (s, 3H). HRMS (ESI-TOF) m/z: [M+H]^+^ calcd for C_15_H_22_N_3_O_2_S, 308.1427; found: 308.1441.

Compounds **11 – 16** were synthesized following a similar procedure for preparing compound **10**.

### (*E*)-1-(4-(dimethylamino)but-2-enoyl)-3-methyl-*N*-(5-methylthiophen-2-yl)azetidine-3-carboxamide (11)

White solid, 43% yield. ^1^H NMR (400 MHz, Methanol-*d*_4_) δ 6.77 – 6.65 (m, 1H), 6.55 (d, *J* = 3.5 Hz, 1H), 6.52 – 6.42 (m, 2H), 4.65 (d, *J* = 8.9 Hz, 1H), 4.35 (d, *J* = 10.6 Hz, 1H), 4.12 (d, *J* = 9.0 Hz, 1H), 3.99 – 3.85 (m, 3H), 2.89 (s, 6H), 2.37 (s, 3H), 1.65 (s, 3H). HRMS (ESI-TOF) m/z: [M+H]^+^ calcd for C_16_H_24_N_3_O_2_S, 322.1584; found: 322.1599.

### (*E*)-2-(4-(dimethylamino)but-2-enoyl)-*N*-(5-methylthiophen-2-yl)-2-azaspiro[3.3]heptane-6-carboxamide (12)

White solid, 16% yield. ^1^H NMR (400 MHz, Methanol-*d*_4_) δ 6.70 – 6.53 (m, 1H), 6.49 – 6.28 (m, 3H), 4.39 – 4.18 (m, 2H), 4.09 – 3.95 (m, 2H), 3.94 – 3.80 (m, 2H), 3.13 – 2.97 (m, 1H), 2.84 (s, 6H), 2.44 (d, *J* = 7.9 Hz, 4H), 2.31 (s, 3H). HRMS (ESI-TOF) m/z: [M+H]^+^ calcd for C_18_H_26_N_3_O_2_S, 348.1740; found: 348.1765.

### (*E*)-4-(dimethylamino)-*N*-(3-((5-methylthiophen-2-yl)amino)-3-oxopropyl)but-2-enamide (13)

White solid, 28% yield. ^1^H NMR (400 MHz, Methanol-*d*_4_) δ 8.15 – 7.88 (m, 1H), 7.71 (s, 1H), 7.49 (m, *J* = 5.8 Hz, 1H), 7.44 – 7.28 (m, 1H), 4.76 – 4.64 (m, 1H), 4.54 – 4.41 (m, 1H), 3.77 – 3.66 (m, 1H), 3.61 – 3.51 (m, 6H), 3.51 – 3.38 (m, 1H), 2.42 (d, *J* = 75.0 Hz, 3H), 1.44 (t, *J* = 7.0 Hz, 2H). HRMS (ESI-TOF) m/z: [M+H]^+^ calcd for C_14_H_22_N_3_O_2_S, 296.1427; found: 296.1452.

### (*E*)-1-(4-(dimethylamino)but-2-enoyl)-*N*-(5-methylthiophen-2-yl)pyrrolidine-3-carboxamide (14)

White solid, 29% yield. ^1^H NMR (400 MHz, Methanol-*d*_4_) δ 6.73 – 6.56 (m, 2H), 6.40 (s, 2H), 3.85 (s, 2H), 3.83 – 3.37 (m, 4H), 3.20 – 3.04 (m, 1H), 2.80 (s, 6H), 2.27 (s, 3H), 2.25 – 1.94 (m, 2H). HRMS (ESI-TOF) m/z: [M+H]^+^ calcd for C_16_H_24_N_3_O_2_S, 322.1584; found: 322.1598.

### (*E*)-1-(4-(dimethylamino)but-2-enoyl)-*N*-(5-methylthiophen-2-yl)piperidine-4-carboxamide (15)

White solid, 43% yield. ^1^H NMR (400 MHz, Methanol-*d*_4_) δ 6.88 (d, *J* = 15.1 Hz, 1H), 6.65 – 6.50 (m, 1H), 6.41 (s, 2H), 4.49 (d, *J* = 12.9 Hz, 1H), 4.07 (d, *J* = 13.5 Hz, 1H), 3.85 (d, *J* = 6.7 Hz, 2H), 3.16 (t, *J* = 12.9 Hz, 1H), 2.82 (s, 7H), 2.59 (t, *J* = 9.8 Hz, 1H), 2.28 (s, 3H), 1.84 (d, *J* = 12.4 Hz, 2H), 1.61 (p, *J* = 13.0 Hz, 2H). HRMS (ESI-TOF) m/z: [M+H]^+^ calcd for C_17_H_26_N_3_O_2_S, 336.1740; found: 336.1766.

### (*E*)-1-(4-(dimethylamino)but-2-enoyl)-*N*-(5-methylthiophen-2-yl)piperidine-2-carboxamide (16)

White solid, 25% yield. ^1^H NMR (400 MHz, Methanol-*d*_4_) δ 7.07 – 6.79 (m, 1H), 6.72 – 6.43 (m, 3H), 5.23 (s, 1H), 4.10 – 3.82 (m, 3H), 3.61 – 3.44 (m, 1H), 2.89 (s, 6H), 2.36 (s, 3H), 2.29 – 2.14 (m, 1H), 1.87 – 1.64 (m, 3H), 1.63 – 1.41 (m, 2H). HRMS (ESI-TOF) m/z: [M+H]^+^ calcd for C_17_H_26_N_3_O_2_S, 336.1740; found: 336.1762.

### *N*-(1-(4-(dimethylamino)but-2-ynoyl)azetidin-3-yl)-5-methylthiophene-2-carboxamide (17)

To a solution of the **I-6** (5.9 mg, 0.03 mmol, 1.0 eq) in DMF (1 mL), HATU (12.5 mg, 0.033 mmol, 1.1 eq), DIEA (35 μL, 0.2 mmol, 2.0 eq) was added followed by 4-(dimethylamino)but-2-ynoic acid (3.8 mg, 0.03 mmol, 1.0 eq). The reaction mixture stirred at rt for 30 mins. Resulting crude mixtures were purified via prep-HPLC to yield titled compound as white solid, 31% yield. ^1^H NMR (400 MHz, Methanol-*d*_4_) δ 7.51 (s, 1H), 6.80 (s, 1H), 4.80 – 4.68 (m, 1H), 4.59 (t, *J* = 8.7 Hz, 1H), 4.40 – 4.32 (m, 1H), 4.31 (s, 2H), 4.29 – 4.23 (m, 1H), 4.13 – 4.04 (m, 1H), 2.97 (s, 6H), 2.48 (s, 3H). HRMS (ESI-TOF) m/z: [M+H]^+^ calcd for C_15_H_20_N_3_O_2_S, 306.1271; found: 306.1289. Compounds **28** was synthesized following the same procedure for preparing compound **17** but with the 4-(dimethylamino)-4-methylpent-2-ynoic acid.

### *N*-(1-(4-(dimethylamino)-4-methylpent-2-ynoyl)azetidin-3-yl)-5-methylthiophene-2-carboxamide (18)

White solid, 28% yield. ^1^H NMR (400 MHz, Methanol-*d*_4_) δ 7.57 – 7.44 (m, 1H), 6.81 (s, 1H), 4.59 (t, *J* = 8.7 Hz, 1H), 4.47 – 4.41 (m, 1H), 4.36 (t, *J* = 9.5 Hz, 1H), 4.26 (dd, *J* = 8.6, 5.7 Hz, 1H), 4.08 (dd, *J* = 10.7, 5.2 Hz, 1H), 2.98 (s, 6H), 2.49 (s, 3H), 1.74 (s, 6H). HRMS (ESI-TOF) m/z: [M+H]^+^ calcd for C_17_H_24_N_3_O_2_S, 334.1584; found: 334.1593.

### *N*-(1-(4-(dimethylamino)-4-methylpent-2-ynoyl)azetidin-3-yl)-5-(methoxymethyl)thiophene-2-carboxamide (19)

To a solution of *tert*-butyl 3-aminoazetidine-1-carboxylate (14.2 mg, 0.1 mmol, 1.0 eq) and 5-(methoxymethyl)thiophene-2-carboxylic acid (17.2 mg, 0.1 mmol, 1.0 eq) in DMF, HATU (41.9 mg, 0.11 mmol, 1.1 eq) and DIEA(0.04 mL, 0.22 mmol, 2.0 eq) were added. The reaction mixture stirred at rt for 60 mins. The resulting crude mixtures were purified via ISCO to yield intermediate. Then the intermediate was dissolved in DCM/TFA (1:1, 1 mL). The reaction mixture was stirred at rt 1 h. After excess TFA was removed, resulting crude mixtures were purified via ISCO to yield intermediate **I-20**. To a solution of **I-20** (6.8 mg, 0.03 mmol, 1.0 eq) and 4-(dimethylamino)-4-methylpent-2-ynoic acid (4.7 mg, 0.03 mmol, 1.0 eq) in DMF, HATU (12.6 mg, 0.033 mmol, 1.1 eq) and DIEA (13 μL, 0.066 mmol, 2.0 eq) were added. The reaction mixture stirred at rt for 60 mins. The resulting crude mixtures were purified via prep-HPLC to yield **(23).** White solid, 31% yield.White solid, 19% yield.^1^H NMR (400 MHz, DMSO-*d*_6_) δ 8.91 (s, 1H), 7.49 (s, 1H), 6.92 (s, 1H), 4.62 – 4.49 (m, 1H), 4.41 (s, 2H), 4.39 – 4.31 (m, 1H), 4.14 – 4.05 (m, 1H), 4.04 – 3.93 (m, 1H), 3.87 – 3.76 (m, 1H), 3.12 (s, 3H), 2.68 (s, 6H), 1.49 (s, 6H). HRMS (ESI-TOF) m/z: [M+H]^+^ calcd for C_18_H_26_N_3_O_3_S, 364.1689; found: 364.1713.

### *N*-(1-(4-(dimethylamino)-4-methylpent-2-ynoyl)azetidin-3-yl)-5-isopropylthiophene-2-carboxamide (20)

White solid, 31% yield. ^1^H NMR (400 MHz, Methanol-*d*_4_) δ 7.55 (s, 1H), 6.88 (s, 1H), 4.83 – 4.71 (m, 1H), 4.61 (t, *J* = 8.3 Hz, 1H), 4.38 (t, *J* = 9.2 Hz, 1H), 4.33 – 4.20 (m, 1H), 4.17 – 4.00 (m, 1H), 3.25 – 3.13 (m, 1H), 2.99 (s, 6H), 1.76 (s, 6H), 1.33 (d, *J* = 6.5 Hz, 6H). HRMS (ESI-TOF) m/z: [M+H]^+^ calcd for C_19_H_28_N_3_O_2_S, 362.1897; found: 362.1916.

### *N*-(1-(4-(dimethylamino)-4-methylpent-2-ynoyl)azetidin-3-yl)-4,5,6,7-tetrahydrobenzo[*b*]thiophene-2-carboxamide (21, MS2159)

White solid, 29% yield. ^1^H NMR (400 MHz, Methanol-*d*4) δ 7.37 (s, 1H), 4.80 – 4.66 (m, 1H), 4.59 (t, *J* = 8.8 Hz, 1H), 4.41 – 4.30 (m, 1H), 4.29 – 4.20 (m, 1H), 4.14 – 3.99 (m, 1H), 2.98 (s, 6H), 2.80 – 2.71 (m, 2H), 2.60 (t, *J* = 6.0 Hz, 2H), 1.88 – 1.77 (m, 4H), 1.75 (s, 6H). ^13^C NMR (101 MHz, Methanol-*d*_4_) δ 164.70, 153.57, 143.66, 137.71, 134.88, 131.09, 86.57, 80.25, 61.72, 58.68, 56.20, 41.39, 39.68, 26.30, 26.06, 24.68, 24.32, 23.70. HRMS (ESI-TOF) m/z: [M+H]^+^ calcd for C_20_H_28_N_3_O_2_S, 374.1897; found: 374.1922.

### *N*-(1-(4-(dimethylamino)-4-methylpent-2-ynoyl)azetidin-3-yl)-4,5,6,7-tetrahydrobenzo[b]thiophene-3-carboxamide (22)

White solid, 41% yield. ^1^H NMR (400 MHz, Methanol-*d*_4_) δ 7.67 (s, 1H), 4.80 – 4.65 (m, 1H), 4.63 – 4.53 (m, 1H), 4.43 – 4.30 (m, 1H), 4.26 – 4.18 (m, 1H), 4.11 – 3.99 (m, 1H), 2.97 (s, 6H), 2.78 – 2.71 (m, 4H), 1.87 – 1.76 (m, 4H), 1.75 (s, 6H). HRMS (ESI-TOF) m/z: [M+H]^+^ calcd for C_20_H_28_N_3_O_2_S, 374.1897; found: 374.1914.

### *N*-(1-(4-(dimethylamino)-4-methylpent-2-ynoyl)azetidin-3-yl)-5,6-dihydro-4*H*-cyclopenta[*b*]thiophene-2-carboxamide (23)

White solid, 33% yield. ^1^H NMR (400 MHz, Methanol-*d*_4_) δ 7.48 (s, 1H), 4.82 – 4.72 (m, 1H), 4.68 – 4.55 (m, 1H), 4.45 – 4.35 (m, 1H), 4.33 – 4.21 (m, 1H), 4.18 – 4.06 (m, 1H), 2.98 (s, 6H), 2.94 (t, *J* = 7.4 Hz, 2H), 2.78 (t, *J* = 7.3 Hz, 2H), 2.48 (p, *J* = 7.3 Hz, 2H), 1.77 (s, 6H). HRMS (ESI-TOF) m/z: [M+H]^+^ calcd for C_19_H_26_N_3_O_2_S, 360.1740; found: 360.1756.

### *N*-(1-(4-(dimethylamino)-4-methylpent-2-ynoyl)azetidin-3-yl)benzo[*b*]thiophene-2-carboxamide (24)

White solid, 26% yield ^1^H NMR (400 MHz, Methanol-*d*_4_) δ 7.99 (s, 1H), 7.90 (t, *J* = 6.6 Hz, 2H), 7.43 (p, *J* = 6.7 Hz, 2H), 4.83 – 4.76 (m, 1H), 4.65 (t, *J* = 8.7 Hz, 1H), 4.42 (t, *J* = 9.5 Hz, 1H), 4.36 – 4.30 (m, 1H), 4.20 – 4.09 (m, 1H), 2.99 (s, 6H), 1.76 (s, 6H). HRMS (ESI-TOF) m/z: [M+H]^+^ calcd for C_20_H_24_N_3_O_2_S, 370.1584; found: 370.1597.

### *tert*-Butyl 2-(azetidin-3-ylcarbamoyl)-6,7-dihydrothieno[3,2-*c*]pyridine-5(4*H*)-carboxylate (I-27)

To a solution of **I-26** (commercially available) (566.6 mg, 2.0 mmol, 1.0 eq) and benzyl 3-aminoazetidine-1-carboxylate (412.4 mg, 2.0 mmol, 1.0 eq) in DMF, HATU(838.6 mg, 2.2 mmol, 1.1 eq) and DIEA(0.7 mL, 4.0 mmol, 2.0 eq) were added. The reaction mixture stirred at rt for 60 mins. The resulting crude mixtures were purified via ISCO to yield intermediate. Then the intermediate was dissolved in MeOH (8 mL) followed by triethylsilane (3 mL). The reaction mixture was stirred at 50 °C overnight in sealed bottle. The resulting crude mixtures were purified via ISCO to yield titled compound as brown oil (101.2 mg, 15% over 2 steps). ^1^H NMR (400 MHz, Methanol-*d*_4_) δ 7.49 (s, 1H), 4.50 – 4.41 (m, 2H), 4.35 – 4.23 (m, 5H), 3.77 – 3.64 (m, 2H), 2.90 – 2.74 (m, 2H), 1.47 (s, 9H).

### 5-(4-aminobutanoyl)-N-(1-(4-(dimethylamino)-4-methylpent-2-ynoyl)azetidin-3-yl)-4,5,6,7-tetrahydrothieno[3,2-c]pyridine-2-carboxamide (I-28)

To a solution of **I-27** (101.2 mg, 0.3 mmol, 1.0 eq) and 4-(dimethylamino)-4-methylpent-2-ynoic acid (46.6 mg, 0.3 mmol, 1.0 eq) in DMF (5 mL), HATU (125.5 mg, 0.33 mmol, 1.1 eq) and DIEA (0.1 mL, 0.6 mmol, 2.0 eq) were added. After stirred at rt for 60 mins, the resulting crude mixtures were purified via ISCO to yield the intermediate. Then the intermeidate was dissolved in DCM/TFA (1:1, 1 mL) followed by stirred at rt for 1 h. After excess TFA was removed, resulting crude mixtures were purified via ISCO to yield titled compounds as brown oil (62.9 mg, 56% over 2 steps). ^1^H NMR (400 MHz, Methanol-*d*_4_) δ 7.59 – 7.49 (m, 1H), 4.34 – 4.22 (m, 3H), 3.62 – 3.51 (m, 2H), 3.24 – 3.17 (m, 3H), 3.16 – 3.07 (m, 3H), 2.98 (s, 3H), 2.92 (s, 3H), 2.87 (s, 3H), 1.74 (s, 3H).

### 5-(9-(3-(6-(1-(2,2-difluorobenzo[d][1,3]dioxol-5-yl)cyclopropane-1-carboxamido)-3-methylpyridin-2-yl)benzamido)nonanoyl)-*N*-(1-(4-(dimethylamino)-4-methylpent-2-ynoyl)azetidin-3-yl)-4,5,6,7-tetrahydrothieno[3,2-*c*]pyridine-2-carboxamide (25, MS2134)

To a solution of **I-28** (7.5 mg, 0.02 mmol, 1.0 eq) and **I-29**^11^ (12.2 mg, 0.02 mmol, 1.0 eq) in DMF, HATU(83.8 mg, 0.022 mmol, 1.1 eq) and DIEA(10 μL, 0.044 mmol, 2.0 eq) were added. The reaction mixture stirred at rt for 60 mins. The resulting crude mixtures were purified via prep-HPLC to yield titled compound as White solid, 24% yield. ^1^H NMR (400 MHz, Methanol-*d*4) δ 8.12 – 8.02 (m, 1H), 7.94 – 7.81 (m, 3H), 7.64 – 7.53 (m, 2H), 7.52 – 7.44 (m, 1H), 7.41 – 7.36 (m, 1H), 7.36 – 7.29 (m, 1H), 7.27 – 7.17 (m, 1H), 4.81 – 4.72 (m, 1H), 4.63 (s, 3H), 4.44 – 4.34 (m, 1H), 4.34 – 4.23 (m, 1H), 4.17 – 4.04 (m, 1H), 3.94 – 3.77 (m, 2H), 3.42 – 3.35 (m, 2H), 3.04 – 2.98 (m, 5H), 2.98 – 2.92 (m, 1H), 2.90 – 2.80 (m, 1H), 2.55 – 2.40 (m, 2H), 2.32 – 2.22 (m, 3H), 1.97 – 1.85 (m, 1H), 1.81 – 1.74 (m, 5H), 1.73 – 1.67 (m, 3H), 1.67 – 1.53 (m, 4H), 1.45 – 1.32 (m, 8H), 1.32 – 1.21 (m, 2H). ^13^C NMR (101 MHz, Methanol-*d*_4_) δ 174.84, 173.99, 169.52, 164.28, 156.83, 153.76, 150.22, 145.33, 144.82, 142.68, 141.79, 141.19, 140.74, 137.27, 136.44, 136.02, 134.56, 134.11, 132.78, 129.58, 128.68, 128.62, 128.35, 128.09, 114.33, 113.68, 111.23, 79.99, 61.25, 58.85, 55.93, 46.63, 44.46, 43.20, 41.57, 41.03, 39.73, 34.50, 34.21, 32.28, 30.36, 30.30, 30.20, 30.18, 27.93, 26.81, 26.47, 26.37, 25.84, 24.98, 19.01, 17.70. HRMS (ESI-TOF) m/z: [M+H]^+^ calcd for C_52_H_60_F_2_N_7_O_7_S, 964.4238; found: 964.4253.

## ASSOCIATED CONTENT

### AUTHOR INFORMATION

#### Author Contributions

Q.W., and X.S. contributed equally to this work.

#### Notes

The authors declare the following competing financial interest(s): J.J. is an equity holder and consultant of Cullgen, Inc., and was a cofounder of Cullgen, Inc., scientific co-founder and scientific advisory board member of Onsero Therapeutics, Inc., and a consultant for EpiCypher, Inc. and Accent Therapeutics, Inc. The Jin laboratory received research funds from Celgene Corporation, Levo Therapeutics, Inc., Cullgen, Inc., and Cullinan Therapeutics, Inc. W.W. has financial interests in Rekindle Therapeutics, where he serves as a co-founder and holds equity. Other authors declare no conflicts of interest.

## ACKNOWLEDGMENTS

This work was supported in part by the endowed professorship from the Icahn School of Medicine at Mount Sinai (to J.J.) and utilized the NMR Spectrometer Systems at Mount Sinai acquired with funding from NIH SIG Grants 1S10OD025132 and 1S10OD028504.

## ABBREVIATIONS

ACN: acetonitrile
Boc: tert-butyloxycarbonyl
DCM: dichloro-methane
DIEA: diisopropylethylamine
DMF: dimethylforma-mide
HATU: O-(7-azabenzotriazol-1-yl)-N,N,N′, N′-tetrame-thyluronium hexafluorophosphate
RT: room temperature
TFA: trifluoroacetic acid
WB: Western blot
WT: wild type

## REFERENCES

1. Bekes, M.; Langley, D. R.; Crews, C. M., PROTAC targeted protein degraders: the past is prologue. Nat. Rev. Drug. Discov. 2022, 21 (3), 181–200.

2. Schreiber, S. L., The Rise of Molecular Glues. Cell 2021, 184 (1), 3–9.

3. Dale, B.; Cheng, M.; Park, K. S.; Kaniskan, H.; Xiong, Y.; Jin, J., Advancing targeted protein degradation for cancer therapy. Nat. Rev. Cancer 2021, 21 (10), 638–654.

4. Hanna, J.; Guerra-Moreno, A.; Ang, J.; Micoogullari, Y., Protein Degradation and the Pathologic Basis of Disease. Am. J. Pathol. 2019, 189 (1), 94–103.

5. Waters, P. J., Degradation of mutant proteins, underlying “loss of function” phenotypes, plays a major role in genetic disease. Curr. Issues Mol. Biol. 2001, 3 (3), 57–65.

6. Chen, Y.; Xue, H.; Jin, J., Applications of protein ubiquitylation and deubiquitylation in drug discovery. J Biol Chem 2024, 300 (5), 107264.

7. Willson, J., DUBTACs for targeted protein stabilization. Nat. Rev. Drug Discov. 2022, 21 (4), 258.

8. Ma, Z.; Zhou, M.; Chen, H.; Shen, Q.; Zhou, J., Deubiquitinase-targeting chimeras (DUBTACs) as a potential paradigm-shifting drug discovery approach. J. Med. Chem. 2025, 68 (7), 6897–6915.

9. Henning, N. J.; Boike, L.; Spradlin, J. N.; Ward, C. C.; Liu, G.; Zhang, E.; Belcher, B. P.; Brittain, S. M.; Hesse, M. J.; Dovala, D.; McGregor, L. M.; Valdez Misiolek, R.; Plasschaert, L. W.; Rowlands, D. J.; Wang, F.; Frank, A. O.; Fuller, D.; Estes, A. R.; Randal, K. L.; Panidapu, A.; McKenna, J. M.; Tallarico, J. A.; Schirle, M.; Nomura, D. K., Deubiquitinase-targeting chimeras for targeted protein stabilization. Nat. Chem.Biol. 2022, 18 (4), 412–421.

10. Liu, J.; Yu, X.; Chen, H.; Kaniskan, H. U.; Xie, L.; Chen, X.; Jin, J.; Wei, W., TF-DUBTACs stabilize tumor suppressor transcription factors. J. Am. Chem. Soc. 2022, 144 (28), 12934–12941.

11. Liu, J.; Hu, X.; Luo, K.; Xiong, Y.; Chen, L.; Wang, Z.; Inuzuka, H.; Qian, C.; Yu, X.; Xie, L.; Muneer, A.; Zhang, D.; Paulo, J. A.; Chen, X.; Jin, J.; Wei, W., USP7-Based Deubiquitinase-Targeting Chimeras Stabilize AMPK. J Am Chem Soc 2024.

12. Deng, Z.; Chen, L.; Qian, C.; Liu, J.; Wu, Q.; Song, X.; Xiong, Y.; Wang, Z.; Hu, X.; Inuzuka, H.; Zhong, Y.; Xiang, Y.; Lin, Y.; Dung Pham, N.; Shi, Y.; Wei, W.; Jin, J., The first-in-class deubiquitinase-targeting chimera stabilizes and activates cGAS. Angew. Chem. Int. Ed. Engl. 2025, 64 (3), e202415168.

13. Wang, Z.; Qian, C.; Xiong, Y.; Zhang, D.; Inuzuka, H.; Zhong, Y.; Xie, L.; Chen, X.; Jin, J.; Wei, W., USP28-based deubiquitinase-targeting chimeras for cancer treatment. J. Am. Chem. Soc. 2025, 147 (16), 13754–13763.

14. Qian, C.; Wang, Z.; Xiong, Y.; Zhang, D.; Zhong, Y.; Inuzuka, H.; Qi, Y.; Xie, L.; Chen, X.; Wei, W.; Jin, J., Harnessing the deubiquitinase USP1 for targeted protein stabilization. J. Am. Chem. Soc. 2025, 147 (17), 14564–14573.

15. Chen, L.; Deng, Z.; Xiong, Y.; Liu, J.; Huang, D.; Wang, J.; Chen, Y.; Inuzuka, H.; Xie, L.; Chen, X.; Jin, J.; Wei, W., Deubiquitinase-targeting chimeras mediated stabilization of tumor suppressive E3 ligase proteins as a strategy for cancer therapy. J. Am. Chem. Soc. 2025, 147 (33), 29875–29883.

16. He, C.; Li, J.; Cui, C.; Xie, J.; Feng, J.; Zhang, B.; Huang, X.; Ma, R.; Zheng, L.; Xu, J.; Yao, H.; Xu, S., Construction of small-molecule deubiquitinase-targeting chimeras to reactivate and stabilize mutant p53 Y220C in vitro and in vivo. Angew. Chem. Int. Ed. Engl. 2026, 65 (3), e18249.

17. Cui, M.; Tang, B.; Yao, T.; Shao, W.; Zheng, Q.; Sun, H.; Hao, H.; Wang, H.; Xu, X., A novel FXR-targeted DUBTAC and its applications in cholestasis therapy. J. Med. Chem. 2026, 69 (7), 7642–7662.

18. Wiener, R.; Zhang, X. B.; Wang, T.; Wolberger, C., The mechanism of OTUB1-mediated inhibition of ubiquitination. Nature 2012, 483 (7391), 618–622.

19. Edelmann, M. J.; Iphofer, A.; Akutsu, M.; Altun, M.; di Gleria, K.; Kramer, H. B.; Fiebiger, E.; Dhe-Paganon, S.; Kessler, B. M., Structural basis and specificity of human otubain 1-mediated deubiquitination. Biochem. J. 2009, 418 (2), 379–390.

20. Song, X.; Wu, Q.; Chen, L.; Inuzuka, H.; Zhong, Y.; Qi, Y.; Lin, Y.; Kabir, M.; Xiong, Y.; Wei, W.; Jin, J., Discovery of new OTUB1 covalent ligands via structure-activity relationship studies for targeted protein stabilization. J. Med. Chem. 2026, 69 (9), 10263–10277.

21. Schapira, M.; Calabrese, M. F.; Bullock, A. N.; Crews, C. M., Targeted protein degradation: expanding the toolbox. Nat. Rev. Drug. Discov. 2019, 18 (12), 949–963.

22. Backus, K. M.; Correia, B. E.; Lum, K. M.; Forli, S.; Horning, B. D.; Gonzalez-Paez, G. E.; Chatterjee, S.; Lanning, B. R.; Teijaro, J. R.; Olson, A. J.; Wolan, D. W.; Cravatt, B. F., Proteome-wide covalent ligand discovery in native biological systems. Nature 2016, 534 (7608), 570–4.

23. Spradlin, J. N.; Zhang, E.; Nomura, D. K., Reimagining Druggability Using Chemoproteomic Platforms. Acc. Chem. Res. 2021, 54 (7), 1801–1813.

24. Park, K. S.; Xiong, Y.; Yim, H.; Velez, J.; Babault, N.; Kumar, P.; Liu, J.; Jin, J., Discovery of the first-in-class G9a/GLP covalent inhibitors. J. Med. Chem. 2022, 65 (15), 10506–10522.

25. De Ioannes, P.; Escalante, C. R.; Aggarwal, A. K., Structures of apo IRF-3 and IRF-7 DNA binding domains: effect of loop L1 on DNA binding. Nucleic Acids Res. 2011, 39 (16), 7300–7.

26. Lukacs, G. L.; Verkman, A. S., CFTR: folding, misfolding and correcting the DeltaF508 conformational defect. Trends Mol. Med. 2012, 18 (2), 81–91.

27. Middleton, P. G.; Mall, M. A.; Drevinek, P.; Lands, L. C.; McKone, E. F.; Polineni, D.; Ramsey, B. W.; Taylor-Cousar, J. L.; Tullis, E.; Vermeulen, F.; Marigowda, G.; McKee, C. M.; Moskowitz, S. M.; Nair, N.; Savage, J.; Simard, C.; Tian, S.; Waltz, D.; Xuan, F.; Rowe, S. M.; Jain, R.; Group, V. X. S., Elexacaftor-Tezacaftor-Ivacaftor for Cystic Fibrosis with a single Phe508del allele. N. Engl. J. Med. 2012, 381 (19), 1809–1819.

28. Wainwright, C. E.; Elborn, J. S.; Ramsey, B. W.; Marigowda, G.; Huang, X.; Cipolli, M.; Colombo, C.; Davies, J. C.; De Boeck, K.; Flume, P. A.; Konstan, M. W.; McColley, S. A.; McCoy, K.; McKone, E. F.; Munck, A.; Ratjen, F.; Rowe, S. M.; Waltz, D.; Boyle, M. P.; Group, T. S.; Group, T. S., Lumacaftor-Ivacaftor in Patients with Cystic Fibrosis Homozygous for Phe508del CFTR. N. Engl. J. Med. 2015, 373 (3), 220–31.

29. Illek, B.; Maurisse, R.; Wahler, L.; Kunzelmann, K.; Fischer, H.; Gruenert, D. C., Cl transport in complemented CF bronchial epithelial cells correlates with CFTR mRNA expression levels. Cell Physiol. Biochem. 2008, 22 (1-4), 57–68.

30. Liu, J.; Chen, H.; Kaniskan, H. U.; Xie, L.; Chen, X.; Jin, J.; Wei, W., TF-PROTACs Enable Targeted Degradation of Transcription Factors. J. Am. Chem. Soc. 2021, 143 (23), 8902–8910.

