## supporting information for "Expanding the Ligandable Chemical Space of OTUB1 through Discovery of a Four-Membered-Ring Recruiter Chemotype"

*Corresponding:

Jian Jin

Wenyi Wei

Yan Xiong

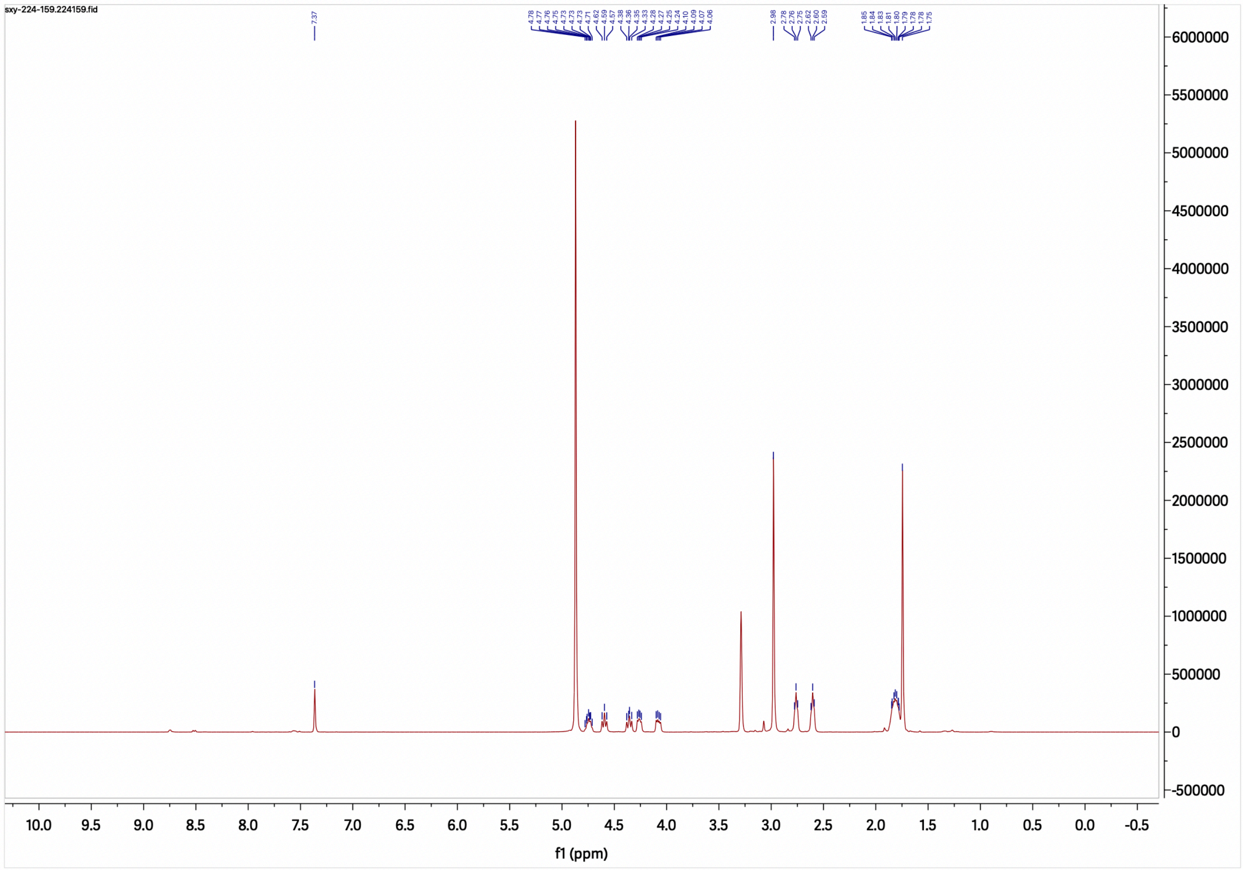

[1]H NMR spectrum of compound 21, MS2159

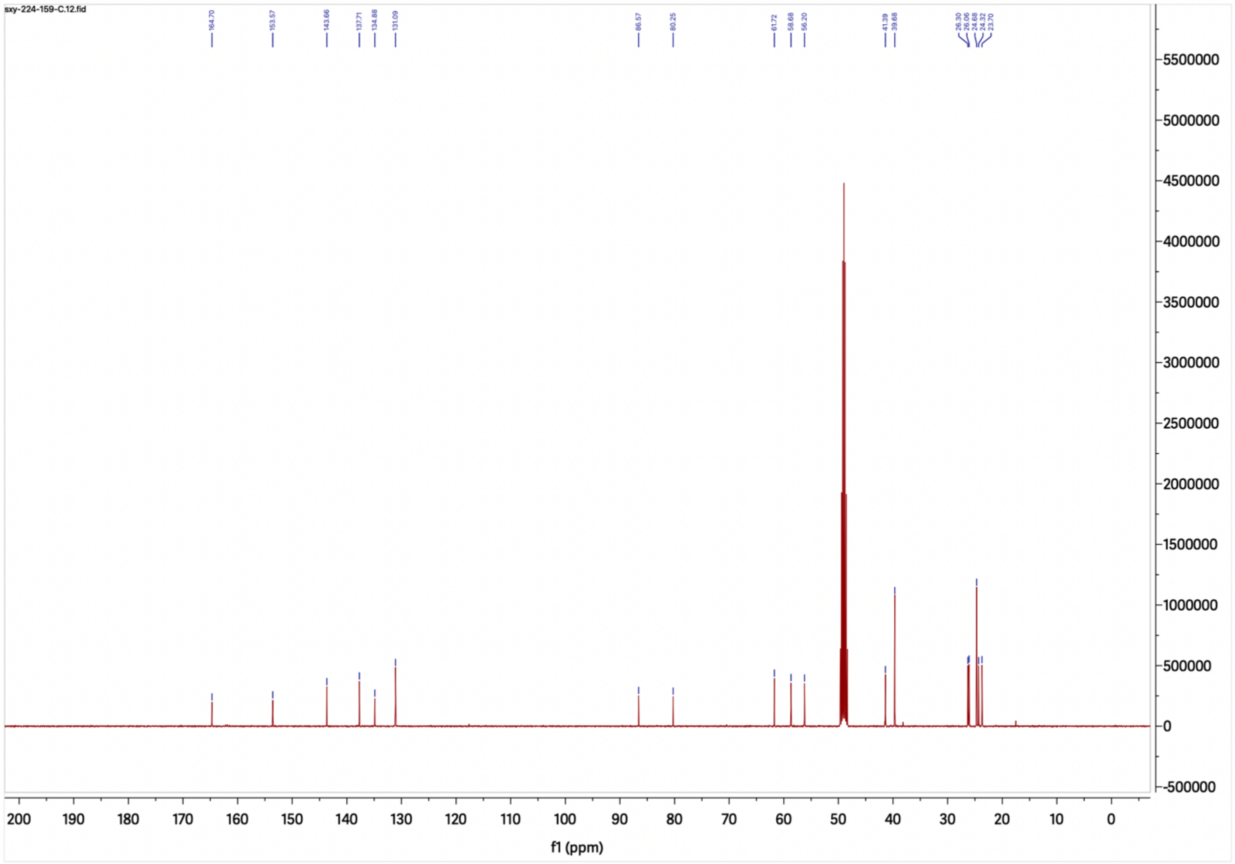


**$^{13}$C NMR spectrum of compound 21, MS2159**

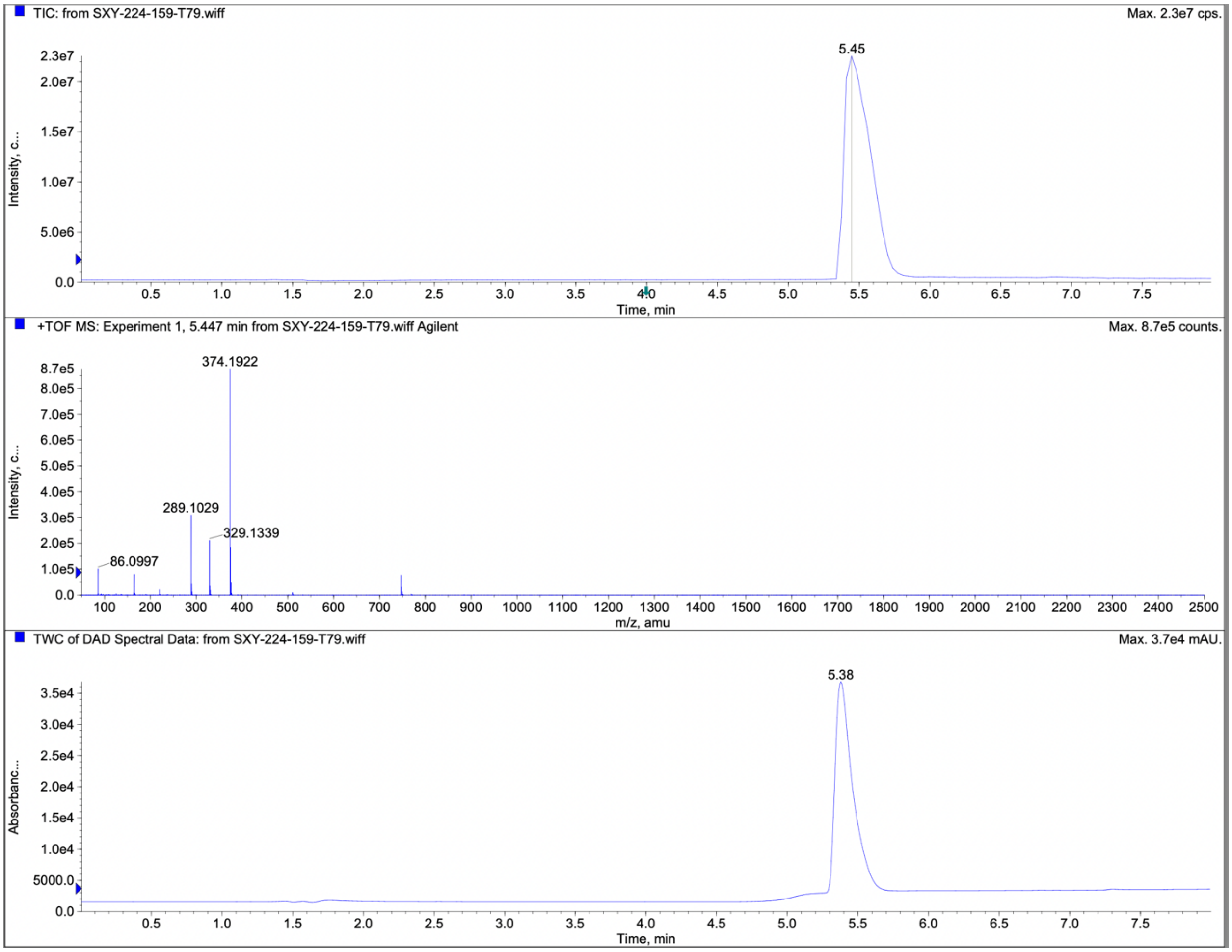


**LC-MS spectra of compound 21, MS2159**

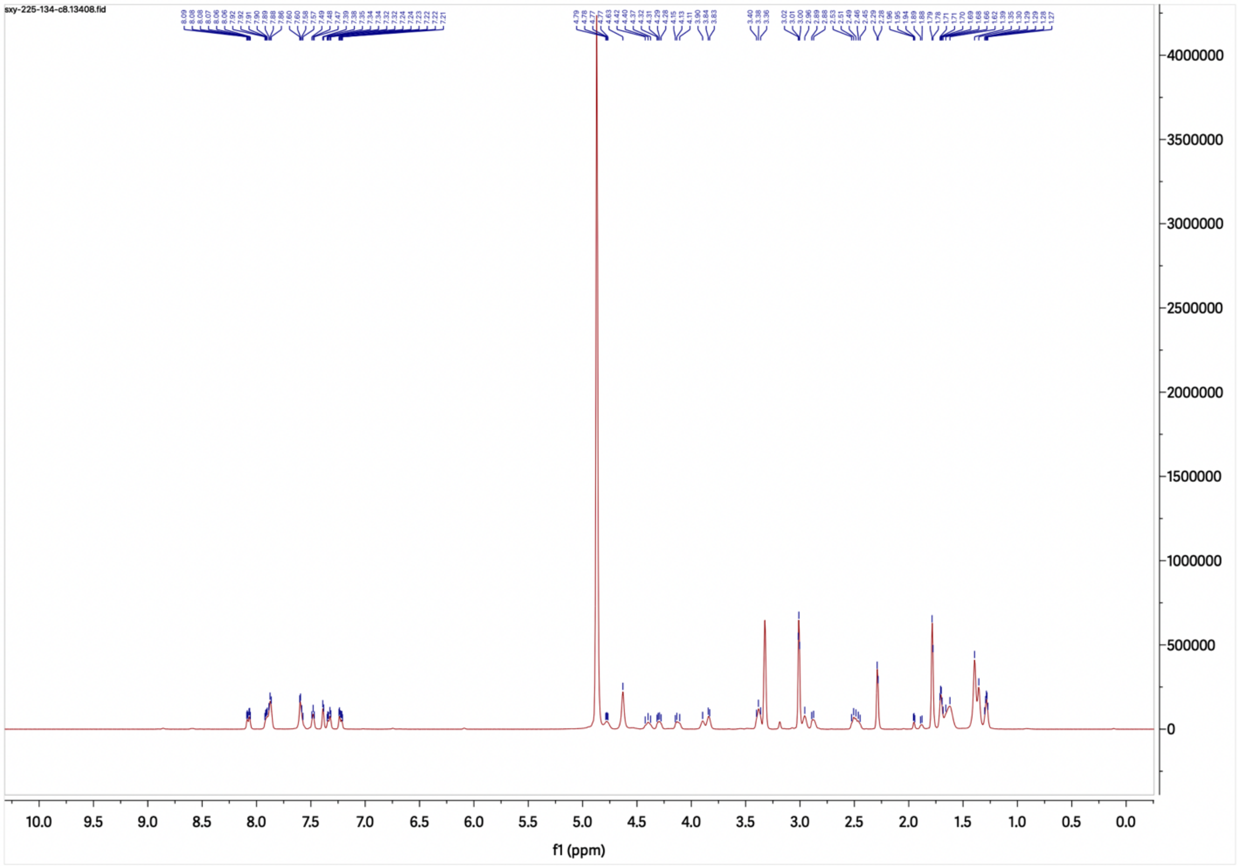


**$^{1}$H NMR spectrum of compound 25, MS2134**

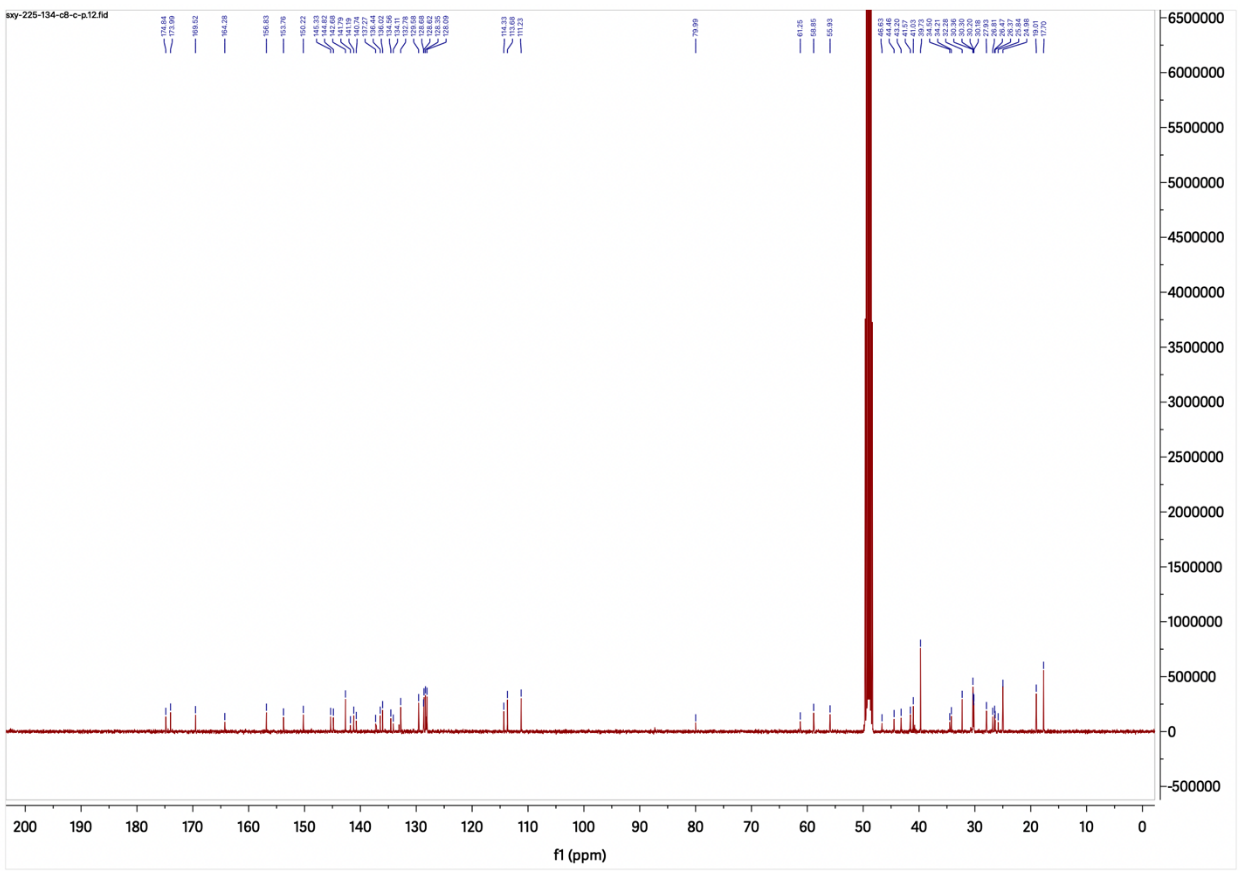

**$^{13}$C NMR spectrum of compound 25, MS2134**

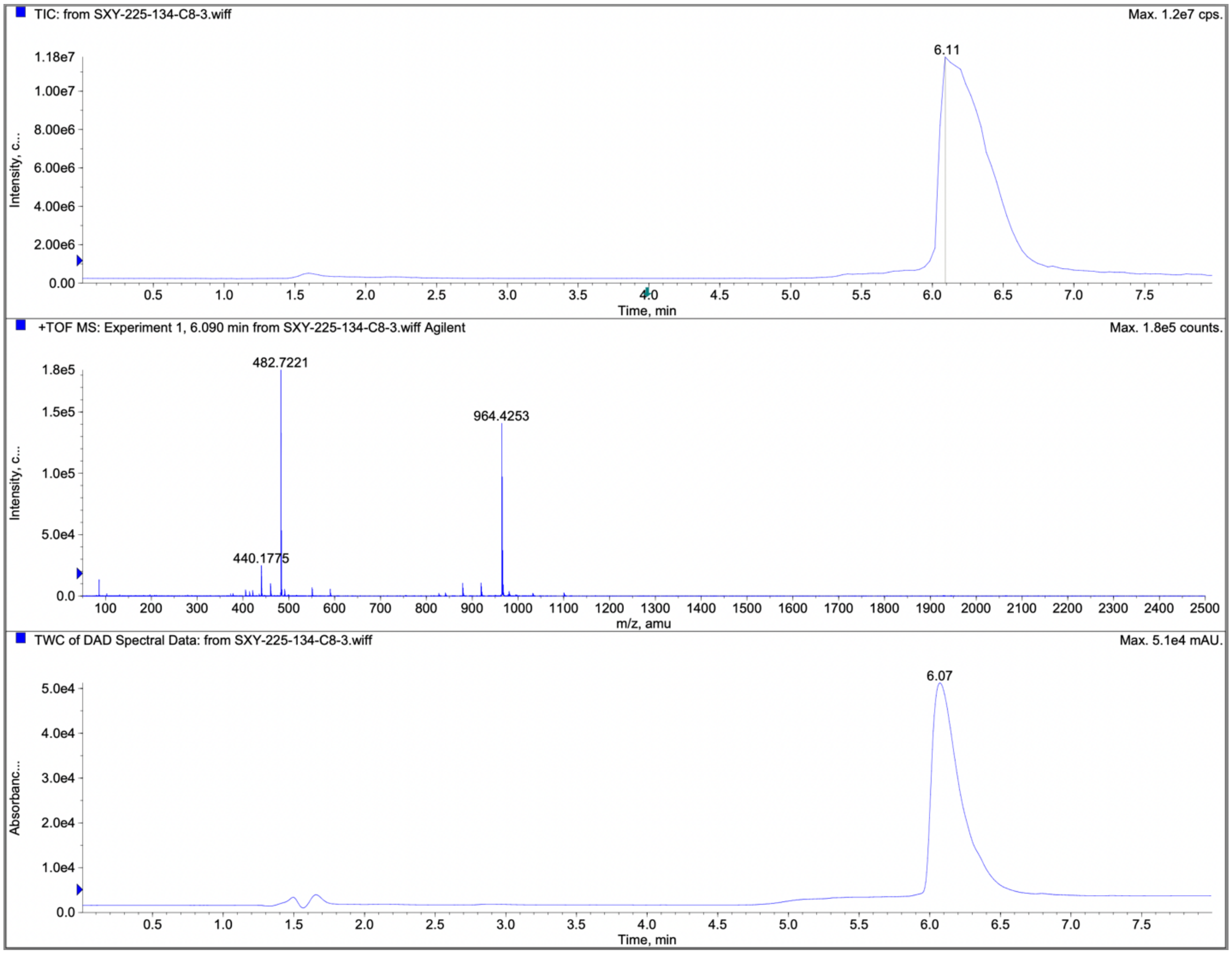


**LC-MS spectra of compound 25, MS2134**
